# Cortical control of innate behavior from subcortical demonstration

**DOI:** 10.1101/2025.02.12.637930

**Authors:** Jason A. Keller, Iljung S. Kwak, Alyssa K. Stark, Marius Pachitariu, Kristin Branson, Joshua T. Dudman

**Affiliations:** Janelia Research Campus, Howard Hughes Medical Institute, Ashburn, VA, USA

## Abstract

Motor control in mammals is traditionally viewed as a hierarchy of descending spinal-targeting pathways, with frontal cortex at the top ^1–3^. Many redundant muscle patterns can solve a given task, and this high dimensionality allows flexibility but poses a problem for efficient learning ^4^. Although a feasible solution invokes subcortical innate motor patterns, or primitives, to reduce the dimensionality of the control problem, how cortex learns to utilize such primitives remains an open question ^5–7^. To address this, we studied cortical and subcortical interactions as head-fixed mice learned contextual control of innate hindlimb extension behavior. Naïve mice performed reactive extensions to turn off a cold air stimulus within seconds and, using predictive cues, learned to avoid the stimulus altogether in tens of trials. Optogenetic inhibition of large areas of rostral cortex completely prevented avoidance behavior, but did not impair hindlimb extensions in reaction to the cold air stimulus. Remarkably, mice covertly learned to avoid the cold stimulus even without any prior experience of successful, cortically-mediated avoidance. These findings support a dynamic, heterarchical model in which the dominant locus of control can change, on the order of seconds, between cortical and subcortical brain areas. We propose that cortex can leverage periods when subcortex predominates as demonstrations, to learn parameterized control of innate behavioral primitives.

## 1. Introduction

Motor control is widely viewed as distributed and hierarchical, with the forebrain and specifically cerebral cortex at the top of the hierarchy ^8,3,9^. However, developmental and lesion studies show that biased connectivity between specific subcortical inputs and outputs enables an innate behavioral repertoire that can be expressed independent of the forebrain ^10–12^. The question of how these innate circuits integrate with forebrain control across several descending motor pathways has a long, unresolved history ^13–16,4,17–22^, and some have suggested a heterarchical arrangement whereby no single node or loop through the environment dominates ^23–28^. This view is not mutually exclusive with hierarchical control, but does imply that a full understanding of forebrain function must be derived from how it learns its effects on the rest of the nervous system and body ^29–31^.

Recordings from frontal cortical areas as well as both acute manipulations and chronic lesions indicate that these areas play a crucial role in (1) dexterous control of movement, particularly limb control in primates and rodents ^32–34^, (2) complex and dynamic feedback control from arbitrary environmental perturbations ^35–43^, and (3) acquisition and modification of stereotyped movements over hundreds to thousands of repetitions ^44–46^, amongst other functions. The role of the forebrain in directing species-typical innate movements, especially over short timescales and/or limited experience remains less clear, but important for understanding how to recover function more efficiently in diseased or injured states. Perhaps the best biological evidence for faster learning processes in cortex comes from work on active avoidance ^47–51^, which highlights frontal cortex as a critical node in a distributed network that enables shuttling movements over freezing to avoid footshock, using sustained and mixed-selective activity. It has been suggested ^50^ that rapidly-learned avoidance responses could be derived from innate behavioral primitives that are known to exist in the hindbrain and spinal cord ^52,53^, but it remains unclear how they could be incorporated into cortical control ^5,7^.

In robots and artificial agents, learning from behavioral primitives has been one of the most successful approaches to efficiently learn robust, flexible, embodied control ^54–56^. In this framework, control networks can learn appropriate parameters to mimic specific movement patterns (motor primitives) following supervised demonstration of a desired primitive. In its most successful form, the primitive demonstration is used to parameterize a dynamical system trajectory. This would appear to have close parallels to proposals that the control of movement by vertebrate neocortex ^35,57,37^ and subcortex ^58,59^ is mediated by low-dimensional dynamical trajectories of neural activity that implement control policies, as well as the observation that motor cortical learning is faster when coupled to an existing neural trajectory ^60,61^. These dynamical trajectories of activity are distributed across cortical and subcortical circuits ^39,46^, but it remains uncertain whether interactions between cortex and subcortex can similarly lead to flexible control of innate behaviors.

A general approach to address these uncertainties includes recording and manipulating neural activity across the neuraxis during a behavior that does not require learning *per se*, yet can be used flexibly in different contexts, a subset of which are likely to require cortical processing. In mice, hindlimb extension stands out in this regard as a fundamental fetal motor primitive ^62^ that is expressed flexibly in the form of rearing and jumping shortly after the ears and eyes open ^10,63^. Furthermore, similar hindlimb extensions are expressed as jumping behavior in adult decerebrate mice ^64^, and can be flexibly controlled using cortical input ^65^. Here we developed a head-fixed task in which mice turn off a cold air stimulus applied to the hindlimbs by vigorously extending them against a counterweight. In this context, naïve mice produce reactive extensions within seconds, consistent with an innate behavioral response, and given the presence of a predictive cue, can learn to produce anticipatory extensions to avoid the cold air in tens of trials. Using 3D kinematic tracking, simultaneous Neuropixels recordings in several brain areas, and acute optogenetic ‘decortication’, we provide evidence for a distributed cortical-subcortical heterarchical control logic in which frontal cortex can learn contextual, predictive motor control from subcortical demonstration of innate motor primitives.

## 2. Results

### 2.1 Head-fixed mice quickly learn predictive control of innate hindlimb extension behavior

Inspired by the remarkable ability of young mice to use vigorous hindlimb extension for escape jumps in various aversive contexts ^63^, we designed a novel head-fixed behavioral paradigm to study the acquired control of innate hindlimb extension. Mice stood on a moveable, counterweighted platform positioned below their hindlimbs, with their forelimbs free to rest upon a curved transparent wall. A controllable stream of cold air was directed downward onto the platform such that hindlimb extension could move the platform downward and increase the air temperature. At a threshold platform displacement the cold air was turned off (Fig. 1a, Suppl. Fig. 1). A motor drives the platform to a starting position and holds it steady to allow a trial structure. Trials were initiated by releasing the platform from a fixed holding position, and a brief auditory cue from a piezo buzzer was provided 3 seconds before the cold air turned on (Fig. 1b). Simultaneous recordings from three cameras, along with refined methods for accurate 3D tracking of keypoints across all limbs, enabled detailed kinematic measurements of extension movements.

**Figure 1:**
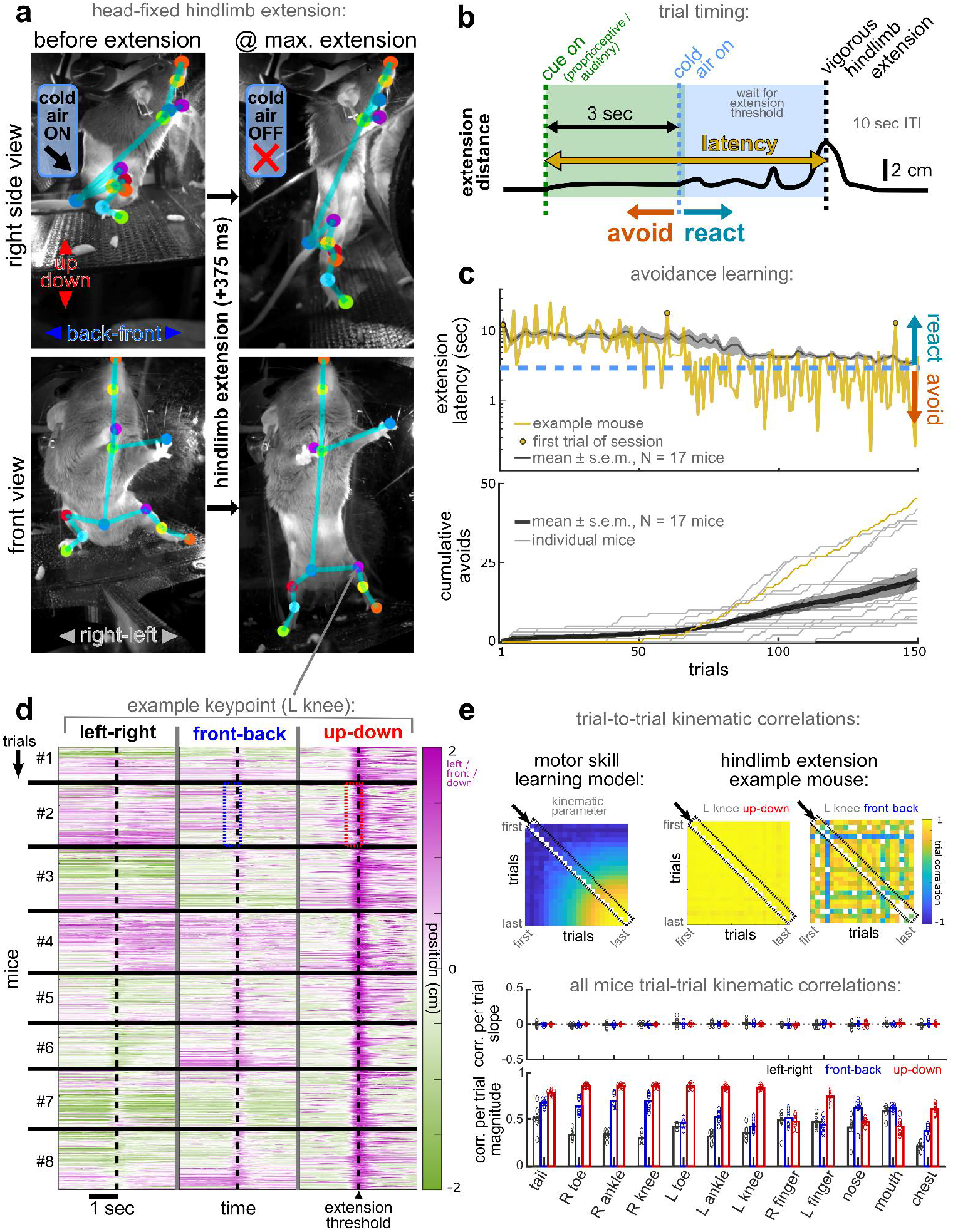
Learning and kinematics during head-fixed hindlimb extension. **(a)** example of head-fixed hindlimb extension behavior from front and side cameras; left frames are 375 ms before extension, right frames are at extension threshold; colored dots are different keypoints, with teal lines added between them for clarity (see also Suppl. Video 1) **(b)** trial timing diagram **(c)** top: latency from cue to hindlimb extension, across trials for an example mouse, and mean ± s.e.m. smoothed over a 5 trial window across all mice; bottom: cumulative avoid trials vs. total trials **(d)** heatmaps of position of left (L) knee keypoint in interval around extension threshold with respect to the median resting position at middle of the inter-trial interval (ITI) across mice, sorted by chronological trials for N=8 mice. Thick black lines separate different mice; grey lines separate columns that reflect the 3 Euclidean dimensions of the keypoint; dotted blue and red boxes show segments for example calculation in (e) **(e)** top left: correlation matrix model for motor skill learning, where trial-to-trial correlations increase over time; top right: examples of up-down and front-back dimension of L knee keypoint over 10-trial bins; black arrow and dotted box show diagonal used for next trial correlations in bottom plots; bottom: mean slopes and magnitudes of the next trial correlations for all keypoints and Euclidean dimensions, for N=8 mice.

First, we find that from the beginning trial mice readily extend their hindlimbs within seconds to turn off the cold air, as expected for innate behavior. Latencies to extend to threshold are initially quite variable but rapidly decrease over the course of 50-100 trials and consistently occur within a few seconds of cold air onset (Fig. 1c; denoted *react* trials). Given the trial structure, mice can also avoid the cold air altogether by extending the platform past the threshold in the time after the platform is released but before cold air turns on. We find that mice generally learn to do so repeatedly within the first 100 trials (Fig. 1c; denoted *avoid* trials). All mice accumulated avoid trials but did not continue to avoid on every trial, instead fluctuating between short bouts of avoid or react trials. Because these bouts do not align across mice, the mean latency per trial remains above the avoid threshold (Fig. 1c, top).

Second, although mice exhibit contextual learning in which they can initiate extensions even in the absence of a direct cold air stimulus (i.e. avoid trials), they do not exhibit systematic changes in kinematic parameters of hindlimb extensions (Fig. 1d-e, Suppl. Fig. 1, Suppl. Video 1). Whereas in motor skill learning (typically over hundreds or thousands of trials) it is typical to observe a slow emergence of stereotyped, correlated kinematic profiles for a movement ^66,44^, we found that movements in the extension direction (‘up-down’, aligned with gravity) are highly correlated across trials from the very start (Fig.1d-e). None of the keypoints show increased correlations in any direction over trials (Fig. 1e), nor is there any systematic increase in bilateral synchronization of hindlimbs (Suppl. Fig. 2). However, we did observe that each mouse adopts an idiosyncratic posture with variance over trials in directions off the axis of gravity (Fig 1d-e).

**Figure 2:**
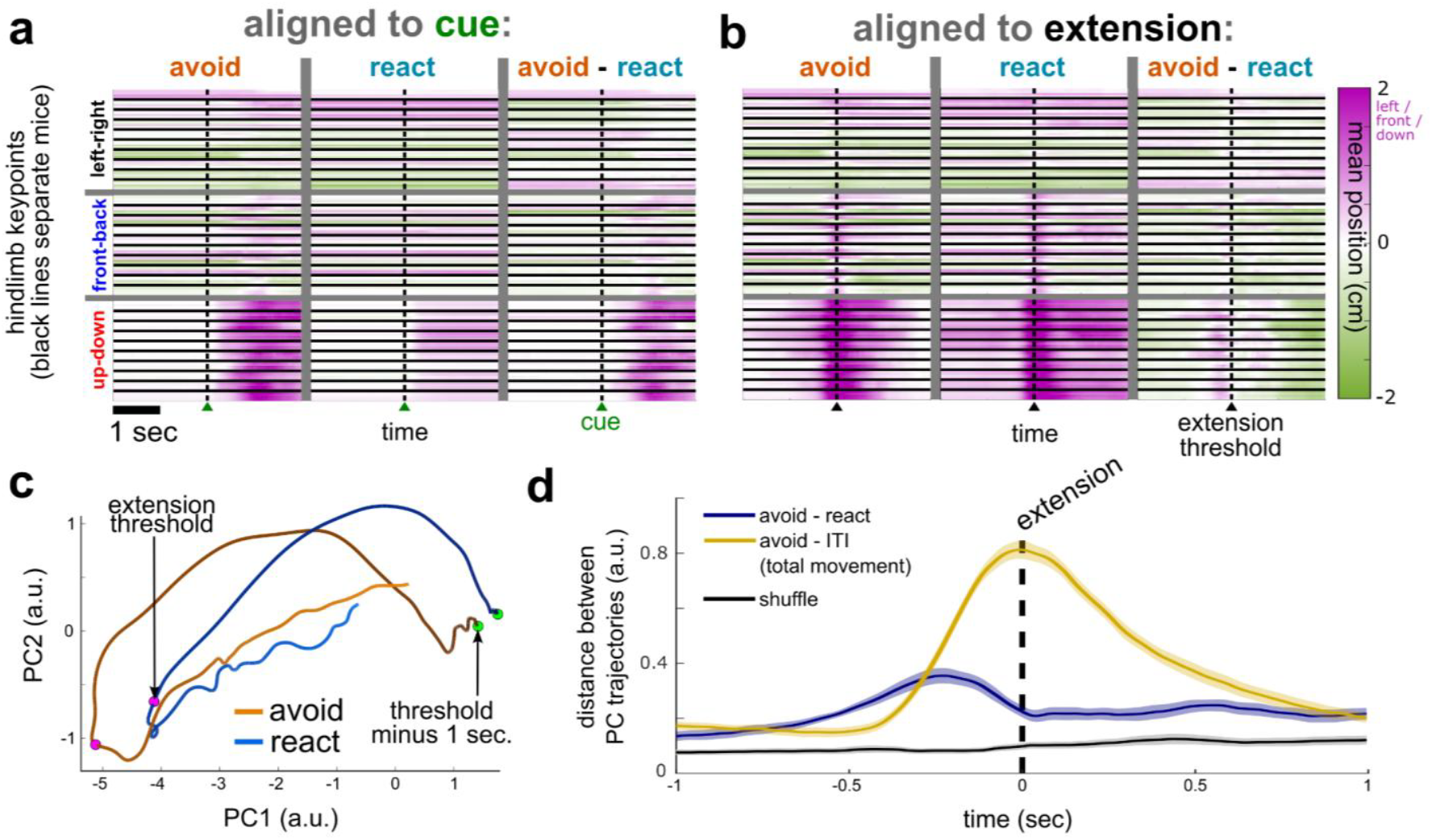
Subtle kinematic differences between avoid and react extensions. **(a)** heatmaps of mean position of all 6 hindlimb keypoints in interval around cue, with respect to the median resting ITI position, during a single session. Thick black lines separate different mice (N=10 mice), and vertical grey lines separate columns that show means on avoid trials, react trials, and the difference between them **(b)** same as (a) but aligned to interval around hindlimb extension **(c)** for an example mouse, trajectories (1 sec. before extension threshold to 1 sec. after) for avoid and react trial means in kinematic state space (first two principal components) based on PCA in interval from 1.5 to 0 sec. before extension threshold **(d)** mean ± s.e.m. (N=10 mice) Euclidean distance across all dimension between avoid and react trial trajectories (purple) in the same interval as (c); gold is positive control distance between avoid extension and mid-ITI interval of same length; black is shuffle negative control distance between two distributions that contain the same number of randomly-selected avoid and react trials, reflecting PCA distance due to random trial fluctuations.

### 2.2 Predictive avoid hindlimb extensions show idiosyncratic posture differences and subtly slower kinematic trajectories

The remarkable kinematic stability of centimeter-scale vigorous extension movements is indicative of an innate movement without clear learned changes in its neural control. However, there are a number of reasons to doubt that similar movements imply similar neural activity, especially in the forebrain. It has long been appreciated ^67,57,37^ and more recently observed in increasing detail ^68–70^ that observed movement kinematics are degenerate with respect to the underlying patterns of neural activity in the forebrain. Moreover, it is unclear how similar a movement would have to be for neural activity to be the same, and thus we sought to examine the kinematics underlying avoid and react trials in greater detail.

The mixture of avoid and react trials within a session and frequently on interleaved trials allowed us to compare the detailed motor (Fig. 2) and neural (Fig. 3-6) correlates of this form of avoidance learning. Direct comparison of mean hindlimb keypoints on avoid versus react trials revealed that they were indeed largely similar (Fig. 2a-b). Alignment to the extension movement revealed reliable differences in the vertical (up-down) dimension, on the order of millimeters over milliseconds, compared to the several centimeters of total movement (Fig. 2b). These differences in vertical movement between avoid and react trials were consistent across mice, showing subtly earlier extension in avoid trials and a tendency to stay extended longer on react trials when cold air was on. Alignment to the platform motor release cue (Fig. 2a) shows that on average all mice extend passively when the platform is released, even on react trials. Furthermore, subtle and idiosyncratic posture differences between trial types in left-right and front-back directions were generally present during the inter-trial interval (ITI, Suppl. Fig. 3), and could reflect a “readiness” for the upcoming cue that predicts its subsequent transformation into an avoid response.

**Figure 3:**
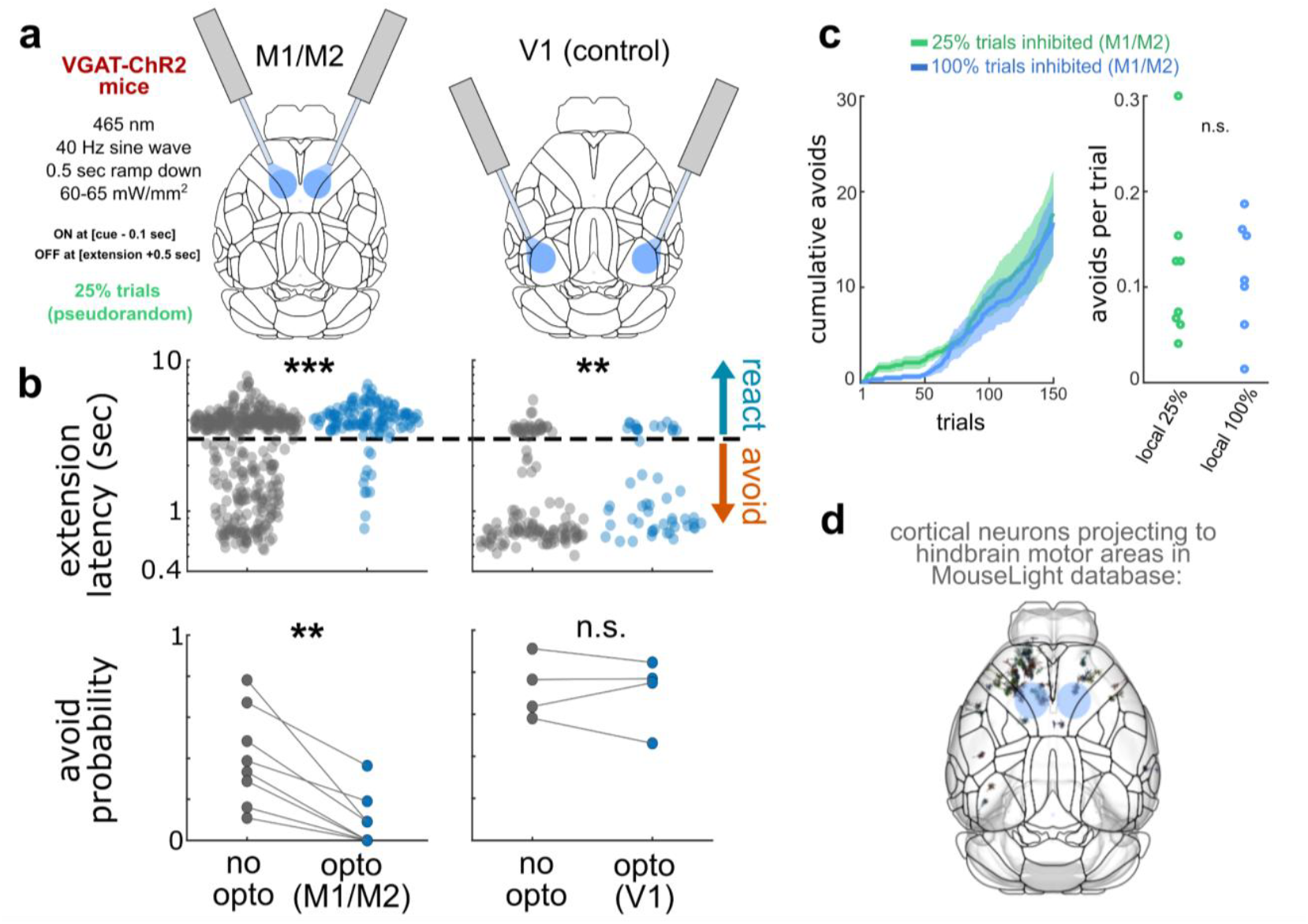
Intermittent local optoinhibition of frontal cortex disrupts avoid but not react hindlimb extensions. **(a)** schematic of bilateral local optogenetic inhibition strategy in frontal cortex (left) versus control visual cortex (right), showing the approximate area of direct inhibition relative to Allen Reference Atlas (https://atlas.brain-map.org/) boundaries **(b)** top, extension latency for each trial with (blue) and without (grey) optoinhibition (frontal/motor M1/M2 cortex, N=8 mice, p=2.12e^-12^ Wilcoxon rank sum test; visual cortex V1, N=4 mice, p=0.006); bottom, avoid probability for each mouse in (b) for trials with (blue) and without (grey) optoinhibition (p=0.008 Wilcoxon signed rank test for M1/M2; p=0.875 for V1) **(c)** cumulative avoid trials vs. total trials for mice with intermittent local optoinhibition of frontal cortex on 25% of trials (teal, mean ± s.e.m. N=8 mice) and different mice with same optoinhibition on 100% of trials (blue, mean ± s.e.m. N=7 mice) **(d)** soma and dendrite locations of cortical neurons that project to hindbrain motor areas implicated in hindlimb extension in the MouseLight database ^75^, with respect to the locally inhibited frontal cortex area (blue circles).

We next combined the 3 Euclidean directions for all 12 keypoints on the body into a 36-dimensional kinematic pose and used principal component analysis (PCA) to capture the dominant kinematic changes across trial types (Fig 2c-d). The top PC in this space accounted for 96.0 ± 1.5% (mean ± s.d.) of movement variance, and was weighted almost exclusively at hindlimb points across all mice, consistent with the majority of reliable movement being in the vertical extension direction (Suppl. Fig. 4). We compared the distance between avoid and react trials in this kinematic state space relative to a control measure of total movement relative to the pose in largely-immobile ITI periods (Fig. 2d). This analysis confirms that the overall kinematic differences between avoid and react trials are small compared to total movement, and that avoid extensions tend to begin a few hundred milliseconds earlier than react extensions, but converge to similar posture at the time of extension threshold.

**Figure 4:**
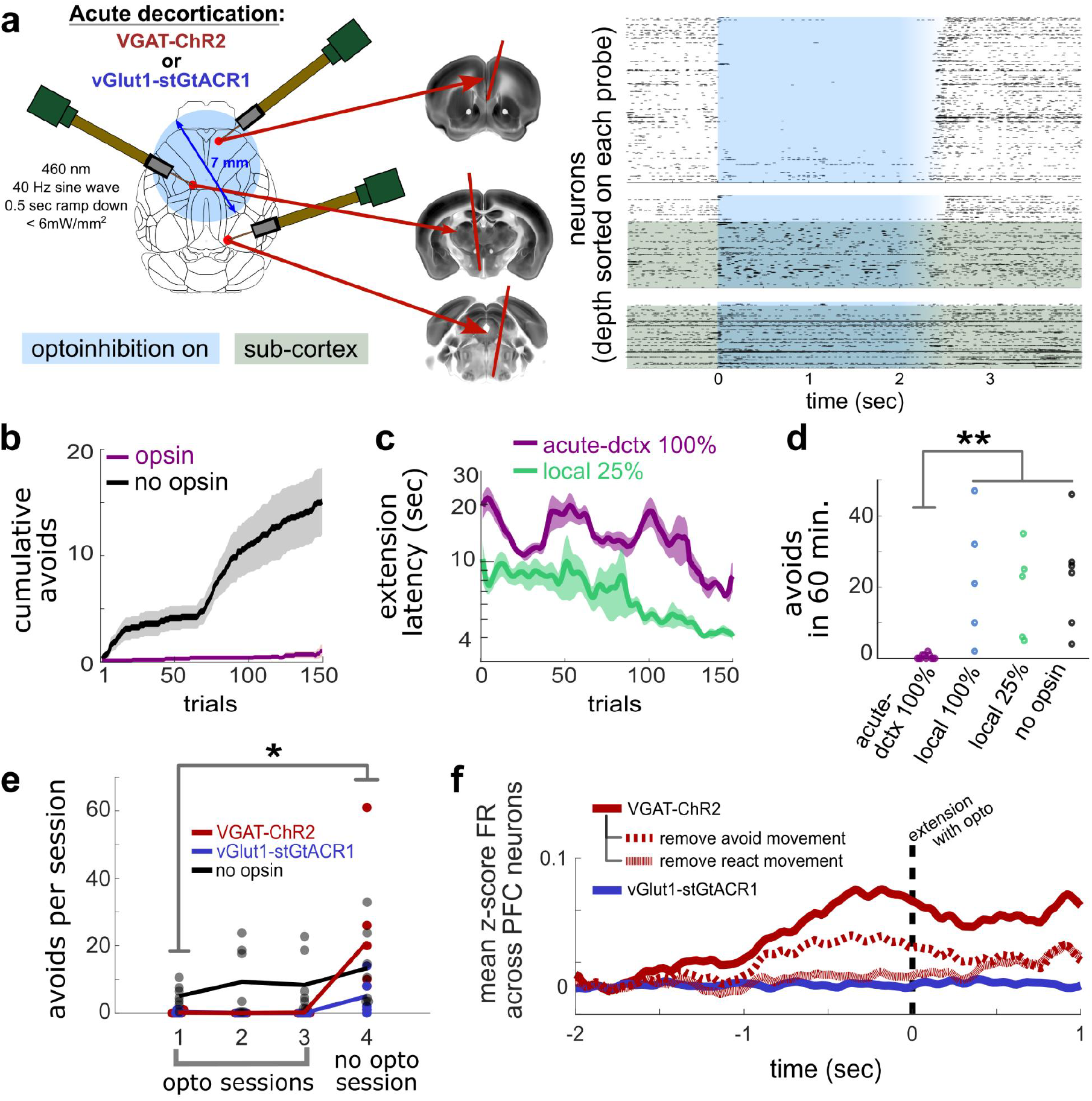
Acute decortication prevents avoid extensions but not learning. **(a)** left, schematic of large-scale cortical optoinhibition (i.e. acute decortication) with simultaneous recordings from 3 Neuropixels probes; right, example recording in vGlut1-stGtACR1 mouse showing near complete acute inhibition over most of dorsal neocortex **(b)** cumulative avoids versus total trials for large-scale optoinhibition (purple, VGAT-ChR2 & vGlut1-stGtACR1 opsin mice combined, N=11; black, wild-type control mice N=7; mean ± s.e.m.) **(c)** extension latency versus total trials for large-scale optoinhibition (purple, same mice as (b); teal, intermittent local M1/M2 inhibition for comparison, N=8 mice; mean ± s.e.m.) **(d)** comparison of total number of avoids over 60 min. of hindlimb extension behavior (each open circle is one mouse; p=0.00021 Kruskal-Wallis test with Dunn-Sidak correction for multiple comparisons) **(e)** avoids per session for VGAT-ChR2 (red, N=6), vGlut1-stGtACR1 (blue, N=5), and wild-type control (grey, N=7) mice; mice with opsins had 3 sessions with optoinhibition on 100% of trials, followed by a session with no optoinhibition; p=0.04 for Wilcoxon rank sum comparison for VGAT-ChR2 mice on session 4 versus no opsin control mice on session 1 **(f)** mean z-scored firing rate (FR) of prefrontal cortex (PFC) neurons during optoinhibited trials, after removal of cold air activity patterns (see Methods: Removing variance), for vGlut1-stGtACR1 mice (N=2) and VGAT-ChR2 mice (N=3, after additional removal of either or react or avoid movement activity patterns).

It is common to invoke “gating” of an evolutionarily conserved caudal brain circuit as the mechanism of innate movement primitives. Indeed, decerebrate mice are able to vigorously extend hindlimbs and escape aversive stimuli ^64^. However, the kinematics on avoid trials are subtly different just prior to extension (Fig. 2d), which could imply a change in the underlying neural control. The subtle change in pre-movement components of the extension on avoid trials is reminiscent of motor preparation characteristic of cortically-controlled movements ^71^. Active avoidance behaviors in other paradigms are also known to depend upon frontal cortex ^48,49^, and thus we next sought to examine the cortical dependence of hindlimb extensions across avoid and react contexts.

### 2.3 Frontal cortex optoinhibition specifically disrupts predictive avoid hindlimb extensions

We used VGAT-ChR2 mice ^72^ to target cortical inhibition to a portion of frontal motor cortex that projects to lumbar spinal cord as well as premotor structures in the hindbrain and brainstem ^73–75^ (Fig. 3a, Suppl. Fig. 5). Inhibiting this part of cortex pseudorandomly on 25% of trials almost completely abolished avoid trials, but mice were still able to execute essentially unchanged react trials with similar latencies and indistinguishable kinematics compared to unperturbed trials (Fig. 3b, Suppl. Fig. 5b, Suppl. Video 2). To control for non-specific perturbation effects, we also examined inhibition of a more caudal region of visual cortex. While this perturbation produced a slight increase in latency on react trials (250.9 msec median difference), there was no change in the probability of avoid trials (Fig. 3b). Finally, releasing optogenetic inhibition 1.5 seconds after trial start (>1 second before cold air onset) yielded normal avoid responses with recovery in <0.5 seconds (Suppl. Fig. 5c), ruling out a nonspecific paralyzation effect.

**Figure 5:**
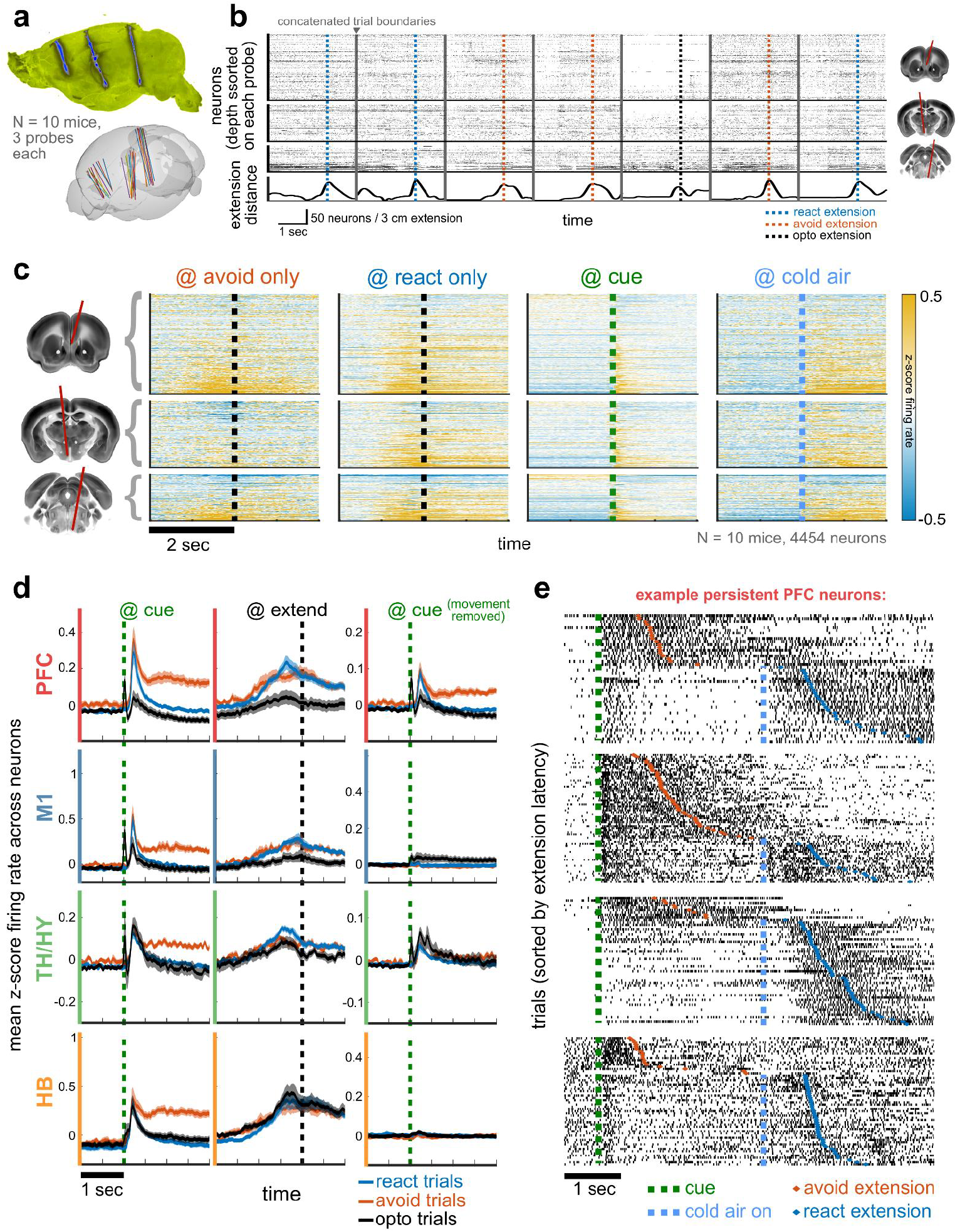
Electrophysiological correlates of avoid and react extensions across forebrain and hindbrain. **(a)** top, max. projection image from light sheet volume of dyed probe tracks (blue) against autofluorescence (yellow); bottom, probe trajectories for all mice (N=10, each mouse has 3 probes of the same color) registered to the Allen Reference Atlas (ARA, https://atlas.brain-map.org/) **(b)** top: example recording from one mouse with acute decortication, showing 7 consecutive trials concatenated around the time of extension; raster shows single spikes, depth-sorted; images on right showing probe locations; bottom: extension distance for example trials **(c)** for the 3 probe locations shown on left, mean z-scored activity pooled across all mice, aligned to avoid/react extension, cue onset, and cold air onset; all heatmaps sorted by modulation magnitude 2 sec before avoid extension relative to baseline ITI baseline **(d)** mean ± s.e.m. z-scored activity for avoid (orange), react (blue), or optoinhibited (black) trials, pooled across all neurons in 4 broad ARA regions (prefrontal cortex, PFC; primary hindlimb motor cortex, M1; thalamus/hypothalamus, TH/HY; hindbrain, HB); left column aligned to cue onset, middle column to extension threshold, right column to cue onset, but after avoid extension activity pattern removal (see Methods: Removing variance) **(e)** trial rasters for example PFC neurons from 4 different mice, sorted by extension latency, showing more persistent activity from cue (dotted green line) to avoid extension (orange diamonds).

The above data indicate that cortical activity is selectively necessary for initiation of avoidance responses, but dispensable for rapid reactive escapes from the cold air stimulus, consistent with previous results of shock avoidance learning ^49^. However, avoidance responses also take several tens of trials to robustly develop (Fig. 1c), indicating that they are learned. A common proposed model for such experience-dependent learning is that robust avoid responses are acquired via reinforcement of cortically-mediated exploratory hindlimb extensions during the pre-cold period. This would suggest that optogenetic inhibition of cortex on all trials could prevent learning of avoid responses. However, we found that focal inhibition of the frontal cortical region described above on 100% of trials only produced a subtle delay in the acquisition of avoidance responses, with mice eventually learning to generate avoid trials at the same rate as those experiencing optoinhibition on 25% of trials (Fig. 3c), similar to the adaptation to optoinhibition recently described in visual cortex ^76^.

The emergence of robust avoidance responses in animals with focal frontal cortical inhibition on every trial implies that there may be a distributed source of descending cortical areas ^77^ that extend outside of our inhibited cortical network, and are sufficient to learn even in the presence of partial, focal inhibition. In line with this possibility, the sampling of cortical projection neurons in the MouseLight database ^75^ reveals that isocortical neurons projecting to brainstem premotor areas implicated in hindlimb extension are found in areas outside the ∼1mm diameter extent of complete local inhibition (Fig. 3d; Suppl. Fig. 5a). By comparison, much prior work on innate behaviors has used large-scale, chronic lesions to produce “decortication” and reveal the critical role of subcortical control. Given that chronic lesions are not applicable to studying dynamic within-session changes, we set out to produce acute, reversible inhibition of the majority of dorsal cortex (“acute optogenetic decortication”) to evaluate whether a distributed cortical network is indeed necessary for mediating the performance and learning of hindlimb extension avoidance responses.

### 2.4 Acute decortication prevents predictive hindlimb extensions, but not learning

To acutely inhibit most of mouse dorsal cortex, we used a powerful LED light source to produce a circular area of inhibition with ∼7 millimeter diameter (∼50x the inhibited surface area typical in optogenetics perturbation experiments, but see ^77^; Fig. 4a, Suppl. Fig. 6). Simultaneous recordings from multiple Neuropixels probes across cortical and subcortical areas verified near complete acute inhibition of cerebral cortex even though the irradiance was lower than for local inhibition (Fig. 4a, Suppl. Fig. 6). Given the large scale and total intensity of light used, we confirmed that heating effects were well within physiological limits and thus effects are attributable to the inhibition of cortical activity (Suppl. Fig. 6d). We used both excitation of inhibitory neurons (VGAT-ChR2 mice) as well as inhibition of excitatory neurons (vGlut1-stGtACR1 mice ^78^) as independent methods to inhibit cortical output, since each has their own caveats ^78–82^.

**Figure 6:**
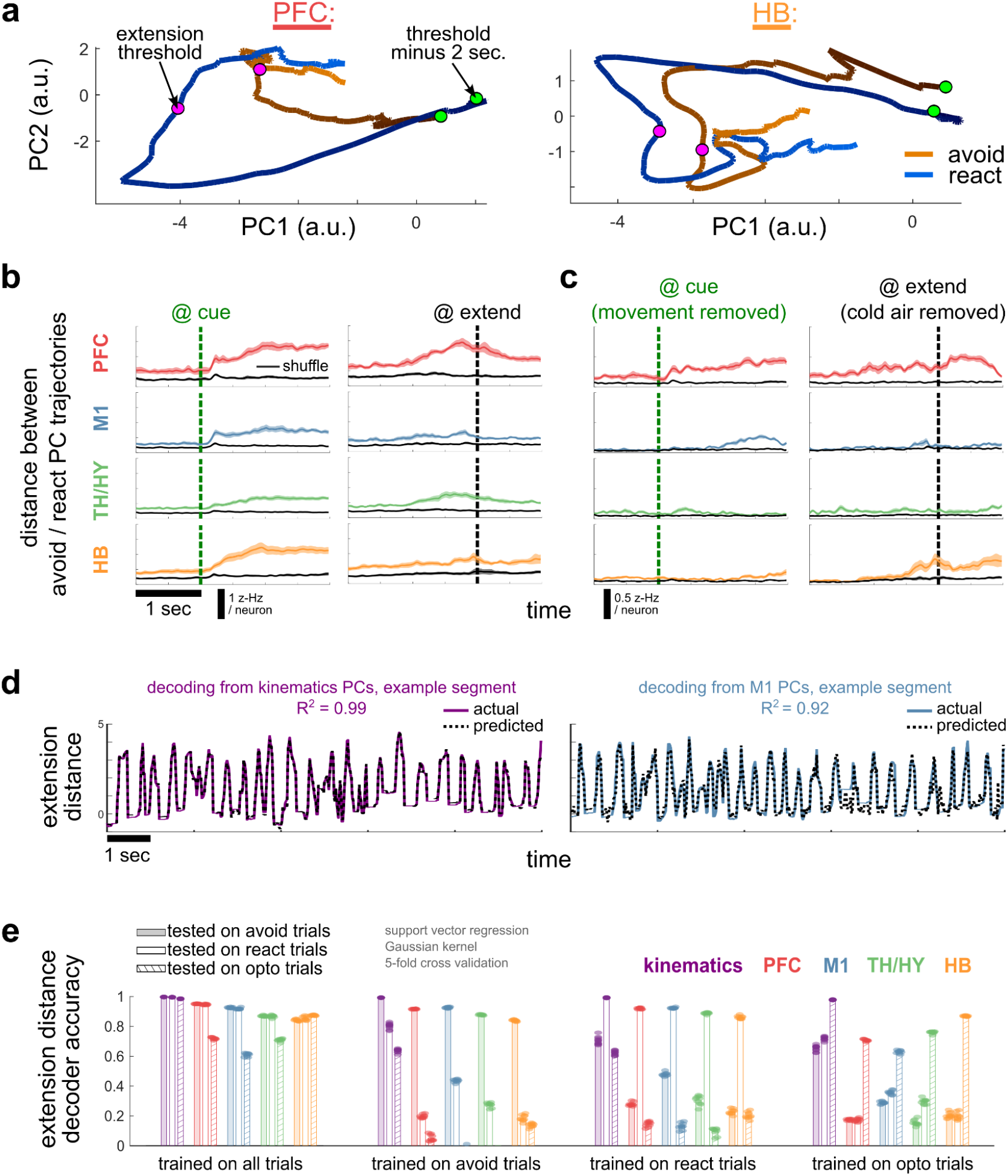
Neural population analyses reveal distributed encoding of avoid versus react extensions. **(a)** example mouse PCA trajectories for avoid and react trial-averaged activity in PFC and HB neurons, from 2 sec. before extension threshold to 1 sec. after **(b)** mean ± s.e.m. PC distances between avoid and react trajectories for different brain areas, aligned to cue (left) and extension (right); black traces are shuffle control reflecting random trial differences **(c)** same as (b) but with removal (see Methods: Removing variance) of avoid movement activity patterns (for cue alignment), or cold air activity patterns (for extension alignment) **(d)** left: example of cross-validation decoding of platform distance from kinematics PCs, using support vector regression; right: same as left but decoding from M1 neural activity PCs **(e)** extension distance decoder accuracy (coefficient of determination, R^2^) from kinematics and four broad brain areas; each dot is mean accuracy across 5 cross-validation folds for a single trial-sampling repeat (N = 10 sampling repeats); accuracies are separated according to trials used for training and test data (all trials, avoid trials, react trials, or optoinhibition trials).

For both methods of acute optogenetic decortication the behavioral effect was the same: essentially complete prevention of avoid trials over 3 sessions even when applied on all trials (Fig. 4b-d). To further control for distraction due to ambient light or non-specific effects of tissue heating we confirmed that control mice lacking opsin-expression showed normal avoidance learning behavior (Fig. 4b,d). These data during both focal inhibition and acute decortication demonstrate that hindlimb extensions can be driven in the absence of cortical input, consistent with recent and classic work indicating that innate behaviors persist following decortication. However, acute decortication did lead to a modest increase in the observed latencies and acceleration of reactive extensions (Fig. 4c, Suppl. Fig. 7c), concomitantly with mice adopting a subtly forward posture and consequent changes in the direction of extensions more forward than down (Suppl. Fig. 7a-b, Suppl. Video 3).

This is consistent with evidence that medial cortical areas are important for trunk stabilization in rats ^83^. Since this perturbation led to increased extension latency (Fig. 4c), we also tested a subset of animals using an extended 10 sec. period between cue and cold air (rather than 3 sec.) to see if they could avoid if given more time. However, all mice still failed to avoid yet responded with similar latency as soon as cold air turned on (Suppl. Fig. 7d), confirming that the cold air stimulus was necessary to motivate hindlimb extension in the absence of cortical output.

After applying acute decortication on every trial for 3 daily sessions, we next released inhibition on the 4^th^ session to test whether acute decortication impaired avoidance learning. To our surprise, we observed an immediate appearance of avoid trials in most VGAT-ChR2 mice as soon as they were released from this large-scale inhibition (Fig. 4e). Thus, mice were able to “covertly” learn to use the predictive cues to elicit avoid trials, even in the absence of any prior cortically-mediated avoidance responses, analogous to the covert learning previously described in songbirds following blockade of forebrain modulation of subcortical premotor structures ^84^. Therefore, mice can learn a cortically-dependent avoidance response even without any trial-and-error experience of driving extensions from the cortex. We propose a simple account: mice can learn directly from a subcortically-mediated “demonstration” of hindlimb extensions in response to cold air. Such learning could, in principle, use information carried via detailed reafference from the periphery or ascending corollaries of subcortical commands ^85–88^. The covertly-learned avoidance responses do not have the same forward posture nor increased acceleration as reactive extensions seen during preceding sessions with acute decortication (Suppl. Fig. 7b-c). This makes learning from detailed reafference less likely, but could still be consistent with subcortical motor corollaries.

To look for evidence of subcortical corollary signals, we examined our simultaneous electrophysiological recordings used to confirm the extent and efficacy of acute decortication (Fig. 4a), particularly the mean modulation of activity aligned to the hindlimb extension for all neurons on the probe targeting prefrontal cortex (PFC; Fig. 4f), where optogenetic light penetration was weakest. There was a clear dissociation in which the cohort of mice showing more robust covert learning (VGAT-ChR2) also exhibited more dorsally-restricted inhibition with more residual movement-aligned activity modulation compared to the cohort with more complete inhibition (vGlut1-stGtACR1). We examined whether this residual activity pattern on optoinhibited trials looked more similar to the movement-aligned activity on avoid trials or non-inhibited react trials (see Methods: Removing variance). If the residual activity was more similar to the activity that drives avoid extensions, it would seem possible that PFC could learn from experience by contributing to optoinhibited react extensions. However, we instead observed that the residual activity pattern is remarkably similar to that on react trials (Fig. 4f), which can be executed with the majority of cortex inhibited, and is thus more consistent with a corollary discharge of subcortical origin.

### 2.5 Sustained and mixed-selective responses to the cue during avoid trials

To this point we have shown that naïve mice escape a cold-air stimulus by evoking an innate, reactive hindlimb extension. Within tens of trials mice can use contextual cues to generate a predictive hindlimb extension that avoids the onset of cold air. The avoid and react movements have subtly different kinematics that were suggestive of changing neural control. Indeed, optogenetic inhibition experiments revealed that a distributed population of frontal cortical populations are critical for generating avoid trials, yet dispensable for reactive extensions to escape cold air. We find that focal optoinhibition can acutely block avoid trials. However, with repeated exposure to focal inhibition, other frontal cortical populations can compensate and restore normal avoidance learning. Complete blockade of avoid extensions requires large-scale optogenetic inhibition approximating an acute decortication, consistent with the possibility that a distributed population of frontal cortical neurons can mediate avoid extensions. Despite a complete blockade of cortically-mediated avoid extensions, avoidance learning can still occur as evidenced by an immediate recovery of avoid trials upon cessation of acute decortication. We propose that such learning can occur via demonstration of reactive extensions mediated by subcortical brain areas.

Together these data make a number of predictions we next sought to confirm with precisely targeted, multi-site Neuropixels recordings from multiple frontal cortical areas as well as midbrain and hindbrain areas associated with hindlimb movements. First, we sought to examine whether there were indeed distributed representations of hindlimb extensions in prefrontal and primary sensorimotor cortex as expected from the optogenetic perturbation experiments. Second, the conjecture that avoidance learning can result from subcortical demonstration implies correlates of subcortically-mediated reactive extension kinematics are present in prefrontal and primary sensorimotor cortex, yet optogenetic inhibition of cortex should revert activity to a more reactive-like activity pattern. Examining these predictions should also put one in position to understand the unique aspects of cortical activity that are critical for avoid trials.

Our core electrophysiology dataset is derived from 10 mice a few days into training, in which there is a mixture of interleaved react and avoid trials (Fig. 5). In each mouse, three Neuropixels probes were implanted to target key brain regions in prefrontal cortex, primary motor cortex, thalamus/hypothalamus, and midbrain-to-hindbrain motor areas implicated in hindlimb extension. An example recording epoch from several concatenated trials reveals large scale coordinated dynamics of activity across cortical and subcortical areas for 3 key trial types: avoid, react, and react during optogenetic inhibition of cortex (Fig. 5b).

The entire dataset consists of 4454 well-isolated single units (neurons). Aligning normalized mean activity of all units to successful avoid extensions and sorting by the magnitude of activity modulation reveals highly distributed correlates of hindlimb extension in subcortical and cortical areas (Fig. 5c, left). Keeping the same sorting across other alignments further reveals substantially overlapping, but also distinct, patterns of activity for react extensions (Fig. 5c, right; Suppl. Fig. 8c). Moreover, many neurons correlated with avoid and/or react hindlimb extensions exhibited robust responses to the trial start cue and the isolated onset of the cold air stimulus; mixed selectivity was thus a prominent feature both of cortical as well as subcortical activity (Fig. 5c, Suppl. Fig. 8).

Using dyed probe tracks to register brains to a standardized anatomical atlas, we matched each recorded unit to a putative brain region in which the neuron was located (see Methods, Fig. 5a). To examine the responses as a function of brain area, we subselected broad anatomical populations of neurons in prefrontal cortex (PFC, red), primary hindlimb sensorimotor cortex (M1, blue), thalamus and hypothalamus (TH/HY; green) and hindbrain (HB, orange; Fig. 5d). Mean z-scored activity in each region revealed robust modulation by trial onset cues and the initiation of a vigorous hindlimb extension for all trial types (Fig. 5d, left). Optogenetic inhibition of cortical activity rendered population activity much closer to the activity observed in react trials, consistent with behavioral results that cortical inhibition reverted mice to a purely reactive mode (Fig. 5d, Suppl. Fig. 9a). The most prominent effect of cortical optogenetic inhibition was the near complete abolishment of a sustained increase in activity following the trial onset cue and associated with avoid trials (Fig. 5d). A portion or even all of this sustained activity could be attributable to hindlimb extensions in the epoch following the trial cue onset. Thus, we next removed the dimensions of population activity associated with hindlimb extension movements to examine residual activity, linearly separable from movement-related activity. This revealed that whereas in HB and M1 essentially all of the sustained, post-trial cue activity on avoid trials was strongly associated with movement, sustained activity in the PFC was preserved (Fig. 5d, right), suggesting the presence of individual neurons in the PFC with sustained activity, whose mean is uniquely linearly separable in avoid trials. Indeed, in all mice we could find such individual neurons, as exemplified in raster plots of example neurons from four mice, having sustained activity from cue to extension in avoid but not react trials (Fig. 5e).

As helpful as it is to examine mean activity modulation around critical behavioral contrasts, there are also limitations to examining mean activity. Models of descending motor control that invoke the importance of low-dimensional dynamics of activity imply that there can be different *patterns* of population activity that produce the same behavioral output ^67,57,37^. Moreover, the fact that the mean modulation of activity during hindlimb extensions is the same for avoid and react extensions in the PFC and HB does not necessarily imply that there are no differences in the pattern of activity within brain areas that can distinguish these trial types. To examine the pattern of activity modulation across the population we next compared activity dynamics projected onto principal components (PCs) - the linear dimensions that capture most variance across the neural population (Fig. 6a). Trajectories of neural activity projected onto the first two PCs revealed a clear separation between avoid and react trials (Fig. 6a). The distance between these dynamical trajectories was largest in the PFC, but avoid and react trial mean trajectories were also well separated (relative to trial shuffled controls) in all recorded areas (Fig. 6b). If we again look at activity linearly separable from that associated with movement, but this time examine the activity projected onto PCs, we again find that much of the differences between avoid and react trials are attributable to movement-related activity in M1, HB and TH/HY. In contrast, well separated trajectories again remain in the PFC, indicating that there is both sustained average modulation of activity and a unique pattern of active cells following the cue, that distinguishes avoid from react trials (Fig. 6c). Given the mixed selectivity observed in all recorded populations (Suppl. Fig. 8), another possibility is that much of this difference at the time of movement is due to the cold air stimulus (which is present during movement in react but not avoid trials). However, when we remove the directions of activity that capture modulation by cold air, we see that a persistent and sustained separation of low dimensional trajectories from cue to extension is uniquely present in PFC (Fig. 6c).

The distances between low dimensional trajectories provide evidence for differences in the pattern of neurons with modulated activity; however, it can also be difficult to quantify such differences. Thus, to complement PCA analyses, we next trained decoders to predict extension distance (derived from independent platform measurement) from neural activity in each recorded brain area. We compared neural decoding performance to that obtained from endpoint kinematics of the animal, which should constitute an upper limit to decoder performance (Fig. 6d). To examine similarities and differences in the extension-related patterns of activity across trial types, we trained decoders either on all trial types and then tested on each trial type separately, or trained on one trial type and computed the cross-validated decoder accuracy (R^2^, see Methods) either on held out trials of the same trial type or trials from the other two types. In all cases, cross-decoding performance declined across trial types, but this decline was modest in the case of kinematics (Fig. 6e) consistent with behavioral results that showed highly similar kinematics across avoid, react, and optoinhibition trials (Fig. 1 & 2, Suppl. Fig 5 & 7). In populations recorded from all brain areas, the decline in decoder performance across trial types was conspicuous and larger than from kinematics (Fig. 6e). This indicates that the highly similar kinematics of hindlimb extensions were mediated by readily separable patterns of active neurons in cortical *and* subcortical areas. Surprisingly, this was even true in HB where a highly conserved pattern activity specific to the innate movement primitive is often assumed. Perhaps most surprisingly, inhibition of cortex did not simply revert HB activity patterns to those of unperturbed react trials, but rather pushed population activity to a new pattern also sufficient to mediate nearly identical hindlimb extensions movements (Fig. 6e). Further consistent with this interpretation, a time-dependent trial-type decoder exhibits similarly good performance in all recorded populations, and even exceeds the ability to use kinematics to distinguish trial types (Suppl. Fig. 9b). This argues that the patterns of neural activity are more distinctive across trial types than the subtle differences in kinematics.

## 3. Discussion

Using 3D kinematic tracking, Neuropixels recordings in several brain areas, and optoinhibition of neocortex, we demonstrate that the generation of kinematically similar hindlimb extensions can switch between dominant cortical and subcortical control on a timescale of seconds. Frontal cortex is dispensable for hindlimb extensions in response to cold air, but necessary for rapidly learned (tens of trials) extensions that precede and thus avoid cold air exposure, using persistent cued activity with mixed-selective representations. Remarkably, however, mice can covertly learn these avoidance responses without any experience of cortically-mediated extensions in our experimental context. Thus, we propose that the cortex “observes” subcortically-mediated hindlimb extension (i.e. “demonstration”) via ascending corollaries and/or reafference to learn arbitrary contexts in which to elicit cortically-driven control of hindlimb extension.

Such shared control across parallel descending motor pathways is not consistent with models of motor control that invoke strict functional hierarchies with a selective emphasis on feed-forward descending pathways carrying motor commands to the periphery. Rather, hindlimb extension in mice appears more consistent with viewing the nervous system as a collection of multiple nested loops through the environment, each having characteristic latencies that operate in concert during the execution of voluntary movements ^89^. The longest “transcortical” loop, reflected in the long-latency reflex component of muscle activity during voluntary movement, is known to be critical for flexible control of movement kinematics in response to external perturbations. When mice are faced with a novel environmental challenge requiring innate hindlimb extension movements, shorter loops can dominate to come up with a more constrained, but “good enough” solution ^6^. In this sense, control is dynamic and heterarchical, and implies that similar learning may occur during development ^31,90^ and in other tasks where initial subcortical control is possible. In line with this view, motor primitives present in both human ^91^ and rodent ^92^ infants have been shown to persist into adulthood.

Where exactly the subcortical demonstration comes from remains an open question. In rodents, many subcortical areas are implicated in hindlimb extension, encoding and escaping cold stimuli, and active avoidance, including but not limited to: spinal interneurons ^93–98^, parabrachial nuclei ^99–101,64^, vestibular nuclei ^102,103^, mesencephalic locomotor areas ^104–107^, periaqueductal grey ^108^, substantia nigra ^109^, zona incerta ^110^, posterior hypothalamus ^111–114^, amygdalar areas ^115,49,116^, and ventral striatum ^117,118^ (but see ^119^). Our examination of activity during acute decortication indicates that subcortical input is critical for learning cortical control, but a full accounting of what subcortical areas are involved and how redundant they are with respect to observed behavior ^120^ will require future studies. Irrespective of the subcortical areas involved, we speculate that the ability of cortex to learn from their demonstration allows a continuum of parameterized cortical control of innate behavioral primitives, from simple gating to tuning all degrees of freedom allowed by the primitive.

It also remains unclear what determines whether or not a mouse will use contextual cortical control to avoid the cold air on a given trial. We show that distributed activity patterns work in concert, with a critical contribution from the transcortical loop through frontal cortex, using persistent and mixed-selective representations in prefrontal areas in particular. The emergence of this persistent PFC activity appears to be a key process in avoidance learning, consistent with a previous study of shock avoidance that also established the importance of amygdalar input ^49^. Our experiments additionally reveal weakly predictive neural activity and posture before an avoid trial even starts, which suggests a potential for “readiness” in the form of internal dynamics, co-contraction impedance control ^121,122^, or even direct tuning of feedback gains at the periphery ^89,123^. Future studies will be required to determine how this readiness leads to persistent PFC activity, and some evidence suggests that neuromodulation could play a key role. Noradrenaline is an intriguing candidate that has previously been shown to modulate the contribution of frontal cortical control in behavior ^124^, but to our knowledge is unknown in this context. Additionally, dopamine release in several brain areas has been implicated in the expression of active avoidance ^125,126,118,127–129^, as well as dynorphin ^130^ and cannabinoids ^118,131^, and all of these may contribute to the learning process as well.

Surprisingly, the avoidance learning process in our experiments did not require cortically-mediated experience, similar to the covert learning previously described in the bird song system ^84^. The residual activity of excitatory neurons deep in prefrontal cortex during acute decortication in VGAT-ChR2 mice appears important for this covert learning, as removing it using vGlut1-stGtACR1 mice prevented learning (Fig. 4). This observation implies an activity-dependent learning mechanism in PFC, but several other mechanisms could be involved. For example, there may be additional plasticity in non-inhibited subcortical areas, non-activity-dependent dendritic plasticity in cortex, or even an enhancement of plasticity at inhibitory synapses across cortical areas, since inhibitory neurons are reliably activated during VGAT-ChR2 cortical inhibition. Teasing apart such mechanisms will be an important goal for future studies.

Covert learning processes challenge conventional reinforcement learning (RL) models in which the expression of a learned action in a specific environmental state necessarily follows from prior experience of that action being executed in that state ^132^. However, a plethora of work in recent years has focused on adding inductive biases to RL models to make them more sample efficient ^133,134^, sometimes combining RL with demonstration ^135^, thus leveraging one of the most successful forms of imitation learning in robotics. Such demonstration learning allows general control policies for primitives to be rapidly deployed in novel contexts or states ^55^. Other recent work has incorporated “action prediction errors” into RL models ^136,137^, and we note that from the point of view of cortex, a dominant subcortical locus of control should induce large errors to learn from, and is consistent with several recent studies demonstrating the ubiquity of motor information across cortex, even in primary sensory areas ^138^. Whatever specific algorithms are used, our work suggests that mammalian nervous systems can use a form of demonstration learning to incorporate inductive biases into cortical processing. This imbues the system with efficiency, flexibility, and robustness, allowing innate behaviors to be executed in a range of contexts beyond the “releasing stimuli” ^14^ with which they co-evolved. Moreover, heterarchical control allows behavior to persist even in the presence of exogenous perturbation (i.e. lesion, inactivation) or endogenous, state-dependent modulation that suppresses initiation in specific learned contexts.

## Methods

### Mice and surgeries

All procedures were in accordance with protocols approved by the Janelia Research Campus Institutional Animal Care and Use Committee. Mice were maintained on a 12:12 reverse light cycle and all experiments were performed during the dark period. VGAT-ChR2-EYFP (JAX#014548), vGlut1-Cre (JAX#037512) x ROSA-stGtACR1 (JAX#033089), and Vglut2-Cre (no-opsin control mice, JAX#16963) mice were used for all experiments. Male and female mice ages 2-6 months were used (no sex differences found in preliminary data analyses, so data is pooled across sexes), single-housed after surgery and given at least three days to recover before starting experiments. Custom RIVETs headplates ^139^ (printed on Formlabs Form3 with Black Resin) were attached to the skull using dental cement (Dentsply Sirona Calibra, Translucent) and superglue (VetBond and Loctite Gel Superglue plus Zipkicker accelerator), leaving as much of the skull as possible exposed for later craniotomies and through skull photostimulation (i.e. “clear skull” preparation ^140^). The skull was protected with silicone elastomer (Smooth On Body Double Fast Set) after surgery and between experiments. Buprenorphine (0.1 mg/kg, subcutaneous injection) was used for postoperative analgesia and Ketoprofen (5 mg/kg, subcutaneous injection) or oral meloxicam (2 mg/kg) mixed with Nutella was used postoperatively for two days. For electrophysiological recordings, a second surgery to make three ∼0.5 mm diameter craniotomies was performed the day before recording started. Craniotomies were drilled using active cooling with a vortex tube (ExAir Mini Cooler) and pointed drill bits (Meisinger HM246-008-HP), and protected with silicone oil and silicone elastomer afterwards.

### Behavior setup and video tracking

The head-fixed hindlimb extension behavioral setup was designed using Onshape CAD and printed with ABS plastic using Stratasys F170 printer. Acrylic walls (McMaster 8560K171 for sides or Grafix K20CP124 for front, laser cut using CAD drawings) were attached to ABS using SciGrip3 acrylic cement. The setup is designed around a lever and counterweight, and uses either common ball bearings (BC Precision 608 Full Ceramic) or optionally an electromagnetic brake (Placid H11-24-1 with tunable constant current PS-24 power supply) as the shaft mechanism. A vortex tube (ExAir 3408 with both hot and cold mufflers to reduce sound) controlled through a solenoid valve (Tameson 3105NC-24VDC) and solid state relay (Numato SSRP2001) provides a cold air stimulus at a ∼45° angle such that it reflects off of the platform floor, and when the floor extends downwards due to hindlimb extension, the temperature increases (Suppl Fig. 1d,e). A linear actuator (Actuonix P16-P) was attached to the counterweight end of the lever to allow pulling the platform back down after a successful jump to start another trial. Three cameras (FLIR BFS-U3-04S2M-CS with Computar A4Z2812CS-MPIR lens; from each lateral side and front) capture synchronized images at 400 Hz and the Exposure Active signal is sent to DAQ streams for alignment with other data, using a Python wrapper around the FLIR API, available at: https://github.com/neurojak/pySpinCapture. Separate camera images were concatenated and stored in MP4 format for later processing using GPU real-time encoding. Infrared LEDs with heatsinks and lenses (Addicore AD476/270/271, 3 on each side) provided illumination at 850nm only (to minimize visual cues available to mice). The platform counterweight consisted of ¼-20 screws and nuts as well as 1-inch pedestal risers (Thorlabs), and the total weight (28 g) was calibrated using gram scale calibration weights placed on the platform, directly under the headplate clamps. The angle of the RIVETS headplate was 35° from horizontal (in the sagittal plane), based on the head angle estimated from video of freely moving vertical jumps. Spatial calibration jigs (RIVETs headplate with extensions orthogonal to the platform floor) were used to align the platform at a consistent height and distance from the point of head fixation. Memory foam (2” thick) cut into small cubes was added to cushion both the bottom and top of platform movement to minimize oscillations. All CAD designs available at https://cad.onshape.com (search: “jumpLever” for rig or “miniRIVETS2” for headplates).

The data acquisition (DAQ) hardware consists of a Dell Precision 5820 PC with NVIDIA Quadro P2000 GPU and PCI-6281 DAQ card (National Instruments, NI) for behavioral control, and a connected PXI chassis (NI-1071 & PCIe-8381) populated with an Imec Neuropixels DAQ card and NI PXIe-6341 for other synchronized analog and digital inputs. Behavior signals were controlled via LabVIEW software (code available at: https://github.com/neurojak/jfcRigControl), which also sends triggers for each camera frame and to SpikeGLX to synchronize with recordings. A LabVIEW state machine detects threshold angular displacements of the platform using input from a calibrated Hall Effect sensor (Littlefuse 55300, Suppl Fig. 1c) with its magnet mounted to the end of the rotation shaft (see CAD design). Once threshold is reached (based on the rolling average of the last 20 samples at 10 kHz), cold air is turned off, and the linear actuator pulls the platform back down and holds it in place for a constant rest period of 10 seconds. To start the next trial, the predictive cue (auditory and proprioceptive information as the linear actuator releases the platform) begins 3 seconds before cold air turns. Because the motor noise cue from the linear actuator does not last until the cold air turns on, additionally a piezo buzzer (4 kHz plus harmonics, CUI CPT-1495C-300) creates an extended auditory cue that terminates either 200 msec after cold air on or after extension threshold is reached in avoid trials, whichever comes first. The extension distance threshold (i.e. Hall Effect sensor voltage) was adjusted to account for mouse size (but held constant across sessions for each mouse), based on the pre-surgery weight at an empirically-based scale of 0.1 V/gram. Daily 20 minute sessions consisted of 70.9 ± 11.4 trials (mean ± s.e.m., N=8 mice over 3-4 sessions), although 9 mice included in Figure 1c had shorter 10 minute daily sessions. For mice used for electrophysiology, craniotomies were performed on the day they first showed reliable avoid responses (day 2 or 3), and recordings were made the following day, in order to ensure a mix of interleaved avoid and react trials.

### Machine vision and kinematics

Twelve keypoints (base of the tail, tip of the nose, tip of bottom jaw center, chest midpoint between shoulders, forepaws 2^nd^ fingertips from pollex, hindpaws middle toe tips, hindlimb ankles front center, hindlimb knees front center) were labeled and tracked using Animal Part Tracker (APT) software (https://github.com/kristinbranson/APT). Three camera views were used to triangulate to 3D space, ensuring that each keypoint could be tracked across at least two cameras in each frame. Mice were shaved caudally from the chest to tail to facilitate keypoint marking, particularly for knee keypoints. Camera parameters were calibrated using OpenCV (https://docs.opencv.org/4.x/d9/d0c/group_calib3d.html) with automatic checkerboard pattern detection, supplemented by manual labelling of stable rig points in all views. Keypoints were labelled independently in all 3 views (but using projected epipolar lines, calculated from calibrated camera parameters to assist labelling of the same point in all views) in a subset of frames that ranged across possible mouse postures on the rig (668 frames total). The tracker, trained using the GRONe algorithm in APT, was subsequently used to automatically track all keypoints. Tracked keypoints were validated using visual inspection of reprojected frames and correlation of tracked hindlimb keypoints with the independent measure of extension distance from the rig angle sensor (0.94 ± 0.03, mean ± s.d., N = 10 mice, Pearson correlation coefficient). Keypoints were rotated from camera space to rig space using SVD to compute a rigid transformation matrix ^141^ from matching stable rig points labelled in camera space and in the CAD model used for rig design. Tracked keypoint positions were subtracted from the overall median across all mice at mid-ITI (which represents a resting pose), and smoothed with a Gaussian window of 50 msec before further analyses. Hindlimb joint angles (Suppl. Fig. 2) were calculated from two keypoint-to-keypoint vectors representing hindlimb segments, as atan2(∥u×v∥,u⋅v), where ∥u×v∥ is the magnitude of the cross product, and u⋅v is the dot product.

### Optoinhibition of cortex

For local optoinhibition, Plexon PlexBright LEDs (465 nm) were used for focal inhibition through the skull in VGAT-ChR2 mice, similar to many previous studies. Patch cords (200/230 um, 0.5 NA) from the LEDs were flat-cleaved and held in place by metal tubes (McMaster-Carr 5560K63) epoxied to a custom 3D-printed piece that maintained spacing across hemispheres and minimized interference with electrodes. The assembly was attached to a Sensapex manipulator using a 6mm rod (ThorLabs) to allow precision targeting of the photostimulation. The end of the patch cords were lowered to touch the skull at the target location (bilateral 1 mm rostral to bregma, 1 mm lateral) and then backed up 200 μm. A 3D-printed plate custom fit to the RIVETs headplates minimized stray photostimulation light reaching the eyes (Suppl Fig. 1b, 6a). The stimulation waveform was a 40 Hz sine wave with 0.5 sec ramp down and average irradiance at the dental cement surface of 60-65 mW/mm^2^. Fibers were centered at the following locations relative to bregma: M1/M2 @ 1 mm rostral (although pilot experiments showed no behavioral difference with 1-2.5 mm placement), 1 mm lateral; V1 @ 3.5 mm caudal, 3 mm lateral. Optoinhibition started just before cue onset on 25% of trials, assigned pseudorandomly (random but not on the first trial or consecutive trials) at the start of the session, and stopped automatically in rare cases if extension threshold was not reached one minute after cold air onset.

For large-scale inhibition of dorsal cortex (i.e. “acute decortication”), two different methods were used, both based on the CoolLED pE-4000 LED light source (Suppl. Fig. 6). The first method used a 3 mm liquid light guide (LLG) coupling from the light source, and refocused its output onto the skull surface using 2-inch relay optics (0.6NA/40 mm FL aspheric condenser, 150 mm FL plano-convex, iris) to create a converging beam with half-max diameter = 6.77 mm (area = 36 mm^2^) at the skull surface, shape measured with a Basler acA4112 (Sony IMX304 sensor) with 2x ThorLabs NE10B ND filters in front to prevent saturation. With the 40 Hz sine wave with 0.5 sec ramp down waveform, average power at the dental cement surface was measured with a ThorLabs S401C sensor to be 180.7 mW total, for irradiance of 5.1 mW/mm^2^. Since cortical heating is a concern ^142,143^, we measured the increase in temperature in the PBS bath at the brain surface using a fast thermistor (TE Measurement Specialties G22K7MCD419 in voltage divider configuration), which was well under physiological limits under worst case illumination conditions (Suppl. Fig. 6d). Because the large optics used in this method can mechanically prevent some desired electrophysiology probe trajectories, we also used a second method in which the LED source was coupled directly to a 2 mm diameter optical fiber patch cord (Doric MFP 1960 μm, 0.5NA), which was held in place by a metal ferrule holder (ThorLabs ADAF4-5) epoxied to a 6mm rod (ThorLabs) to allow precision placement above the skull with minimal electrode interference. Calibrated headplate landmarks were used to center this diverging beam such that it created a 7mm diameter spot on the skull surface, to match the beam in the LLG with optics method above as close as possible. Total power was also calibrated to match the first method, with irradiance at the dental cement surface of 5-5.1 mW/mm^2^. In both methods the beam was aligned such that its caudal edge was 2.5-3.0 mm caudal to bregma, such that very little of visual cortex and none of superior colliculus received direct illumination. For vGlut1-stGtACR1 mice, pilot experiments at irradiance = 5 mW/mm^2^ caused overt time-locked orofacial movements, and thus irradiance was titrated down until no visible movement was induced, to 0.9-1 mW/mm^2^, which still provided powerful cortical inhibition (Fig. 4a).

### Electrophysiology and probe localization

Probe tracks were planned using Pinpoint ^144^ and Neuropixels Trajectory Explorer (https://github.com/petersaj/neuropixels_trajectory_explorer ) software. Neuropixels 1.0 probes ^145^ with tips coated in 1uL CM-DiI for track visualization were mounted on micromanipulators (Sensapex uMp on custom-machined stands), and lowered to their target locations plus 100μm at 5μm/sec, then slowly backed up to target. Recording started after waiting at least 10 minutes for drift stabilization.

SpikeGLX and CatGT (https://billkarsh.github.io/SpikeGLX) were used to record and post process raw voltage data (CatGT parameters: -apfilter=butter,12,300,10000, -gblcar, -gfix=0.4,0.1,0.02), and Kilosort4 ^146^ plus Phy2 (https://github.com/cortex-lab/phy) was used for spike sorting and curation (see details below). Asynchronous clock rates on different headstages and data acquisition cards were pre-calibrated over 1+ hour test runs in SpikeGLX, ensuring sub-millisecond synchronization between behavioral and electrophysiological signals over our shorter recording durations. Ground wires from different probes were tied together (and REF connected to GND) and connected to a silver wire (after soaking in bleach) and placed into the same bath of sterile PBS that contacted the brain surface. After recording, animals were perfused with 4% PFA and whole brains were cleared and imaged using LifeCanvas SmartBatch+ delipidation and a SmartSPIM light sheet microscope. Autofluorescence in the deep red channel was used to register brains to the Allen Reference Atlas using brainreg ^147^ software, and the red channel CM-DiI signal was used to map the probe trajectory and assign each electrode an anatomical location. We chose 3 standard probe trajectories based on previous studies implicating these areas in control of hindlimb extension and avoidance behavior. Probe entry locations and trajectories relative to bregma were:

<u>Probe 1) Prefrontal cortex (right side):</u> craniotomy at 2.8mm rostral, 1.1mm lateral; medial-lateral (M-L) angle 17°, rostral-caudal angle = 19°, standard Bank 0 IMRO table, insertion depth 3.6mm

<u>Probe 2) Hindlimb motor cortex plus underlying thalamus and hypothalamus (left side):</u> craniotomy at 0.4mm caudal, 1.0mm lateral; medial-lateral (M-L) angle 9°, rostral-caudal angle = 1°, long column Bank0/1 IMRO table, insertion depth 5.7mm

<u>Probe 3) Visual cortex, cuneiform nucleus, pontine & medullary reticular areas (right side):</u> craniotomy at 4.1mm caudal, 1.5mm lateral; medial-lateral (M-L) angle 9°, rostral-caudal angle = 11°, long column Bank0/1 IMRO table, insertion depth 5.8mm

### Spike sorting

Raw electrode voltages were preprocessed using CatGT for bandpass filtering, common average referencing, and correction of photostimulation artifacts (see parameters above). Putative single units were isolated using Kilosort4 spike sorting and Phy2 manual curation. Default Kilosort4 parameters were used, except for template detection threshold (=15), spike detection threshold (=13), batch size (=0.5 sec of samples), and number of blocks (=1 for rigid drift correction). We used conservative detection thresholds to ensure units had robust amplitude signals that are stable across any potential movement artifacts. ‘Good’ clusters from Kilosort were curated using Phy2 by discarding clusters with either (1) less than 20 spikes over the session, (2) spatial extent > 200*μ*m, (3) lack of refractory period in the autocorrelogram, (4) spike amplitude that clearly drifted out of the detection threshold over the recording, or (5) spike waveform or amplitude distribution not clearly distinguishable from the noise floor. Occasional merges and splits of clusters were performed based on spike amplitude distributions and principal components of spike waveforms. Ten experimental sessions (each from a different mouse, all with 3 simultaneous probes targeted as described above) yielded 4454 total units across all mice (135.0 ± 52.1 units per probe, mean ± s.d.).

### Data analysis and statistics

All analyses were done in MATLAB using custom code, available with data and other manuscript information at: https://tinyurl.com/Neurojak2025). Nonparametric statistics were used unless otherwise stated, with relevant p-values and tests given in figure legends. Extension distance was calculated based on angle sensor calibration data (Suppl. Fig 1c) along with the arclength of the platform movement, using a radius of the median hindpaw resting position. Velocity and acceleration were calculated as the first and second derivatives of extension distance. Avoid extensions were detected using MATLAB findpeaks() (MinPeakHeight = 1cm) on the extension distance trace in a 3 sec window prior to early cue termination. The distance peak nearest to cue termination was marked as the avoid extension time.

To compute mean electrophysiological responses across trials, spike times were binned at 20ms and z-scored, then smoothed with a half-Gaussian window of 5 bins / 100 msec for further analysis. To compare avoid and react trials, only trials without optoinhibition, as well as react trials without subthreshold extension attempts during the avoid interval (defined at extension distance peaks > 1cm), were included, thus ruling out any trial type similarities due to inhibition effects or unsuccessful avoid attempts. Data were aligned across trials at cue onset (when platform release motor command is sent, although it takes ∼120 msec to process the command, overcome inertia, and start moving), hindlimb extension (angle sensor threshold crossings when the cold air was turned off), or midway through the 10-second intertrial interval (mid-ITI).

### Trial sampling and shuffle controls

Where data were compared across avoid, react, and/or optoinhibited trials, the same number of trials was used for each, by taking the minimum number of trials for each mouse, across conditions, and using random sampling from the other conditions with greater numbers of trials. The random sampling procedure was repeated 10 times to control for trial sampling bias. Shuffle controls for random trial-to-trial differences were computed by creating two shuffled trial distributions for each mouse, using the same number of trials and repeats as used for the comparison data.

### Principal component analyses

We used PCA to reduce the dimensionality of patterns across all kinematics keypoints (Fig, 2, Suppl. Fig. 4) or neurons (Fig. 6, Suppl. Fig 9), and quantify differences between trial types. We used mean data across trials for both kinematics and electrophysiology to create covariance matrices from which PCs were computed using singular value decomposition (SVD). For kinematics data, we used a cue-alignment window from cue onset to 1.5 sec after, and an extension-alignment window from extension threshold reached to 1.5 sec before, and computed PCs separately for each mouse. For neural data, we aligned PCs across mice using an approach similar to a recent study ^50^, by pooling neurons across mice before performing PCA. We used alignment windows to both cue and extension (2 sec before and after), for both avoid and react trial means. Thus, the aligned PCA space represents the linear dimensions that best capture variance between avoid and react trials around the time of both cue and extension. We used SVD to compute PC coefficients, and then QR decomposition to re-orthogonalize the coefficients for a particular mouse, using only the top ten PCs. This alignment de-emphasizes mean responses that differ across mice due to idiosyncratic differences, and allows combining trials across mice for comparison of dynamics and decoding across brain regions, using an equal number of PCs (rather than varying numbers of neurons recorded in each mouse and brain region), at the expense of slightly less variance explained. Neurons were pooled in broad regions such that all recordings had at least 20 neurons in each region for each mouse.

Distances between PC trajectories were calculated at each timepoint as the Euclidean distance across all dimensions between respective projected points in PCA space, normalized by the square root of the number of dimensions used for projection. Where PCA trajectories were compared across trial types, shuffle control distances between PC trajectories were computed using two shuffled trial distributions for each mouse as described above. The two trial-shuffled means were projected onto the same PCs used for the main comparison, and the Euclidean distance between them was computed in the same way. For clarity, PC distances were smoothed using a 500 msec half-Gaussian window.

### Removing variance related to movement or cold air

When comparing neural activity across trial types, in addition to comparing all dimensions of the data, for some analyses we also removed the influence of the potentially confounding dimensions of movement or cold air, for further comparison (Fig. 4f, Fig. 5d, Fig. 6c). To do so, we removed the variance associated with activity patterns recorded during specific time windows. For data aligned to the time of extension, we used a window around cold air stimulus onset (0.5 sec before to 0.2 sec after) to remove activity patterns associated with cold air, which occurs during react but not avoid trials. For data aligned to cue onset, we used a window before extension threshold on avoid trials (1.0 to 0 seconds before) to remove movement variance during the cue period that occurs in avoid but not react trials. For each condition, neural activity from the relevant temporal windows was averaged across trials to generate a condition template. Then we used SVD on these templates to identify linear dimensions in the neural data explaining more than 99.9% of condition variance, and the remaining dimensions were used to construct a projection matrix that removes the variance associated with that condition from the original neural activity space. Removing template conditions from data in PCA space (Fig. 6c) was similar, except that we constructed the projection matrix with respect to the top 9 of 10 dimensions in the aligned subspace across mice (instead of with respect to dimensions accounting for 99.9% of variance in neural data space).

### Platform distance decoders

To compare more equally across brain areas, we decoded from the top ten aligned PCs (smoothed using a 500 msec half-Gaussian window) for each mouse and brain region, instead of directly from neural data having different numbers of neurons in each brain region and across mice. We used support vector regression (MATLAB fitrsvm() function with Gaussian kernel, BoxConstraint=1, KernelScale=1), decoding platform distance in a window [-1.5 to 0.0] seconds relative to extension threshold reached for each trial. Five-fold cross-validation was used to compute the mean coefficient of determination (R^2^) fit of actual and predicted platform distance.

### Trial type classifiers

Trial type was classified (Suppl. Fig 9b) using a support vector machine classifier (MATLAB fitcsvm() function with Gaussian kernel, BoxConstraint=1, KernelScale=1), also from the top ten aligned PCs (smoothed using a 500 msec half-Gaussian window) in each brain region. The classifier was trained separately for every 20 ms bin, using data aligned at cue onset. Five-fold cross-validation was used to compute mean ± s.e.m (across trial samples) classification accuracy at each time bin.

## Acknowledgements

We thank members of the Dudman lab for project feedback; Juan Gallego for comments on the manuscript; Ben Foster and Morgan Clark for help with histology; Michael DeSantis for help with light sheet imaging; Bill Karsh and Jennifer Colonell for help with SpikeGLX and electrophysiology analysis; members of Janelia Project Technical Resources led by Gudrun Ihrke for assistance with labeling keypoints (A.K.S., Emily Tenshaw) and imaging (Christina Christoforou). This work was supported by the Howard Hughes Medical Institute (HHMI).

This article is subject to HHMI’s Open Access to Publications policy. HHMI lab heads and Project Team Leaders have previously granted a nonexclusive CC BY 4.0 license to the public and a sublicensable license to HHMI in their research articles. Pursuant to those licenses, the author-accepted manuscript of this article can be made freely available under a CC BY 4.0 license immediately upon publication.

## Contributions

J.A.K. designed experiments, collected and analyzed data, and wrote the manuscript. S.K., A.K.S., and K.B. assisted with kinematic tracking. M.P. assisted with spike sorting and provided early access to Kilosort4. J.T.D. designed experiments, analyzed data, and wrote the manuscript.

## Supplementary Figures

**Supplementary Figure 1:**
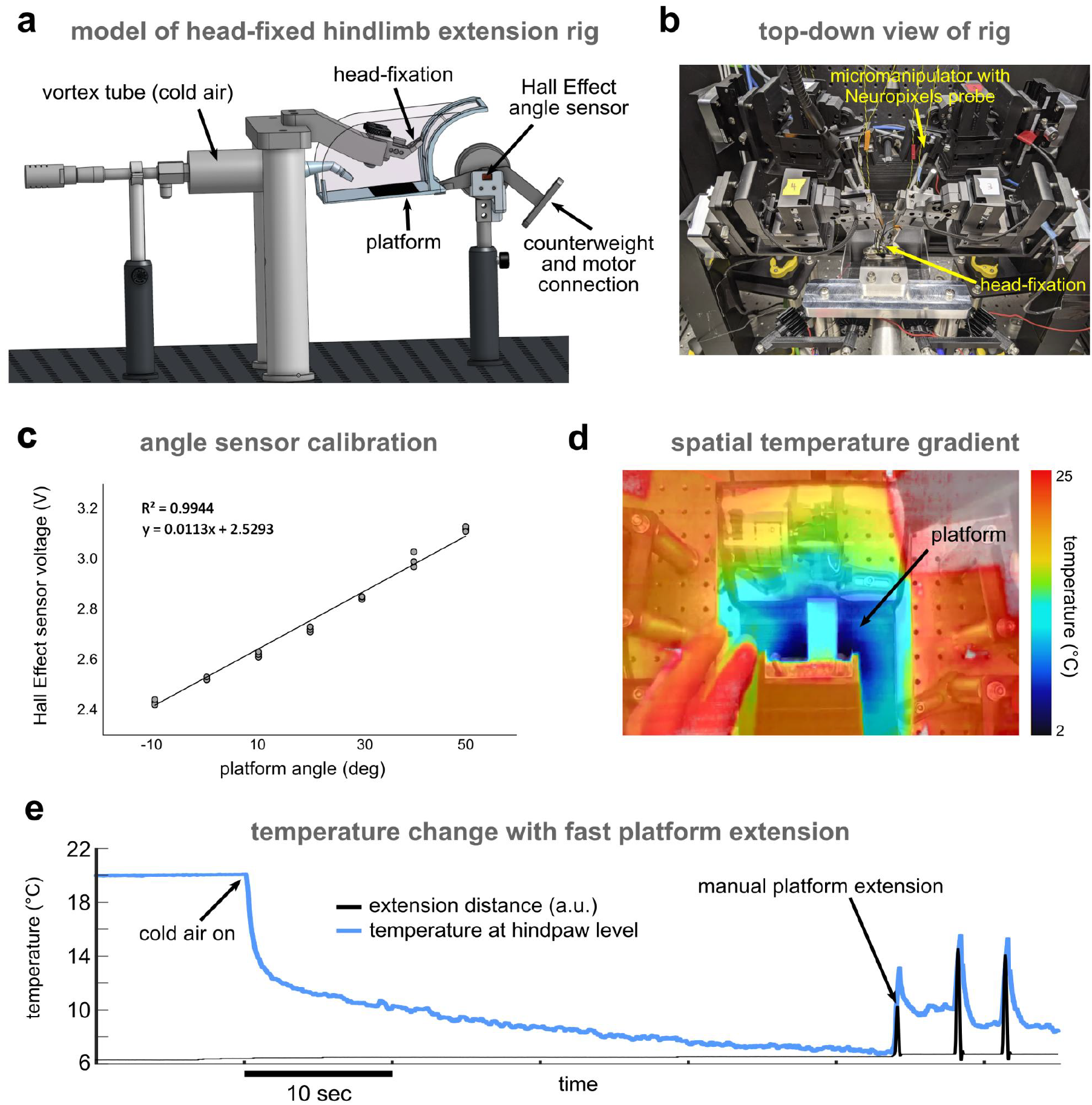
Behavior setup details. **(a)** annotated partial CAD design of behavioral rig **(b)** photo of rig from top back view showing manipulators for electrophysiology and optoinhibition plus custom light shield around head fixation **(c)** Hall effect angle sensor approximate voltage linearity with extension angle/distance **(d)** thermal image of temperature gradient on the mouse platform **(e)** thermistor measurement of rapid temperature decrease at cold air onset, then increase due to platform extension.

**Supplementary Figure 2:**
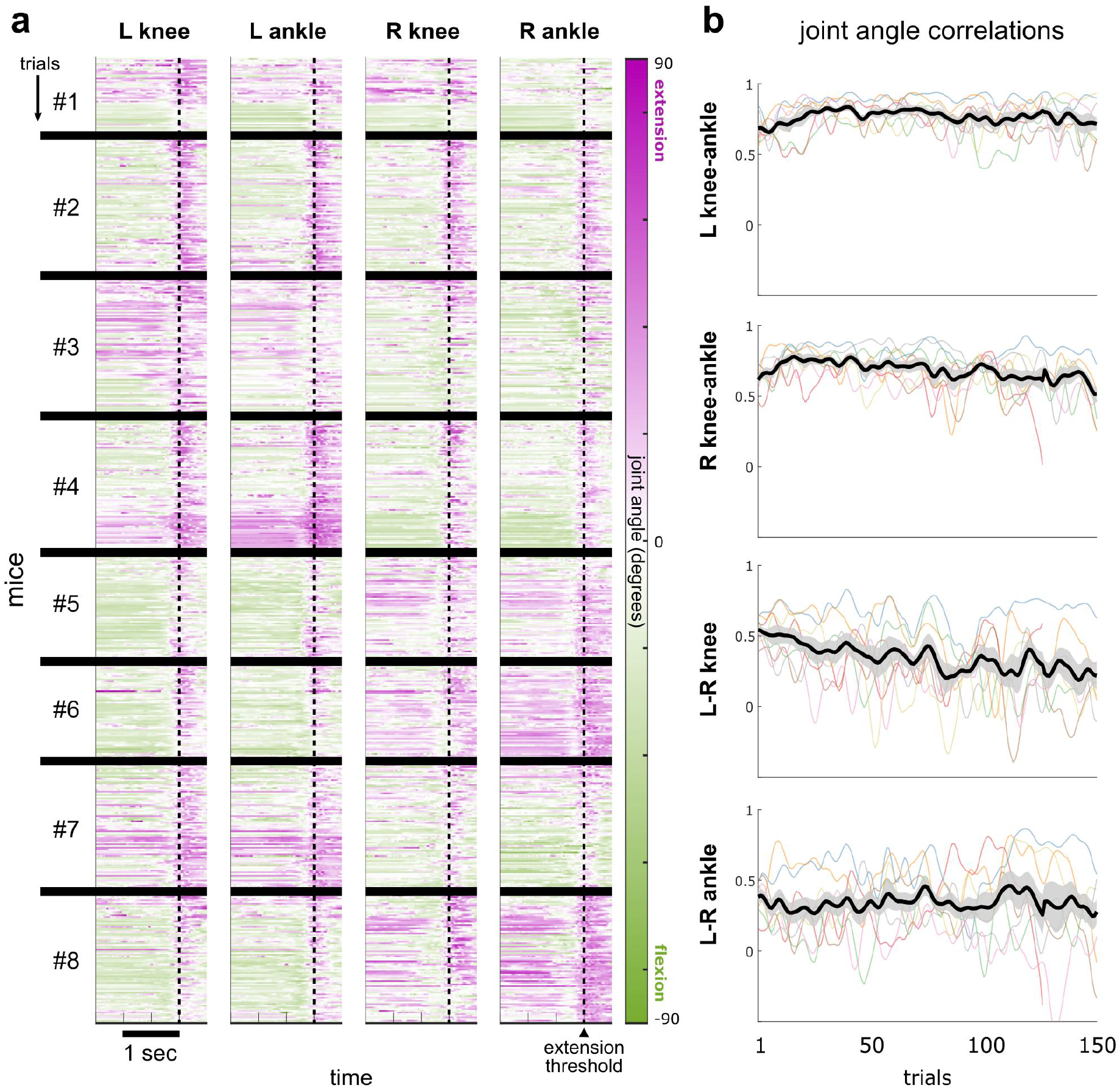
No systematic differences in bilaterality of hindlimb extensions over learning. **(a)** heatmaps of knee and ankle angles in interval around extension threshold, sorted by chronological trials and relative to grand mean angle across all mice and trials, N=8 mice; thick black lines separate different mice; knee angle was approximated based on the angle between vectors (1) from base of tail to knee keypoints, and (2) from knee to ankle keypoints; ankle angle was approximated based on the angle between vectors (1) from knee to ankle keypoints, and (2) from ankle to toe keypoints **(b)** hindlimb angle correlations (Pearson correlation coefficient) over trials, smoothed over a 10 trial Gaussian window (thin colored lines are individual mice, thick black line/shading is mean ± s.e.m. for N=8 mice).

**Supplementary Figure 3:**
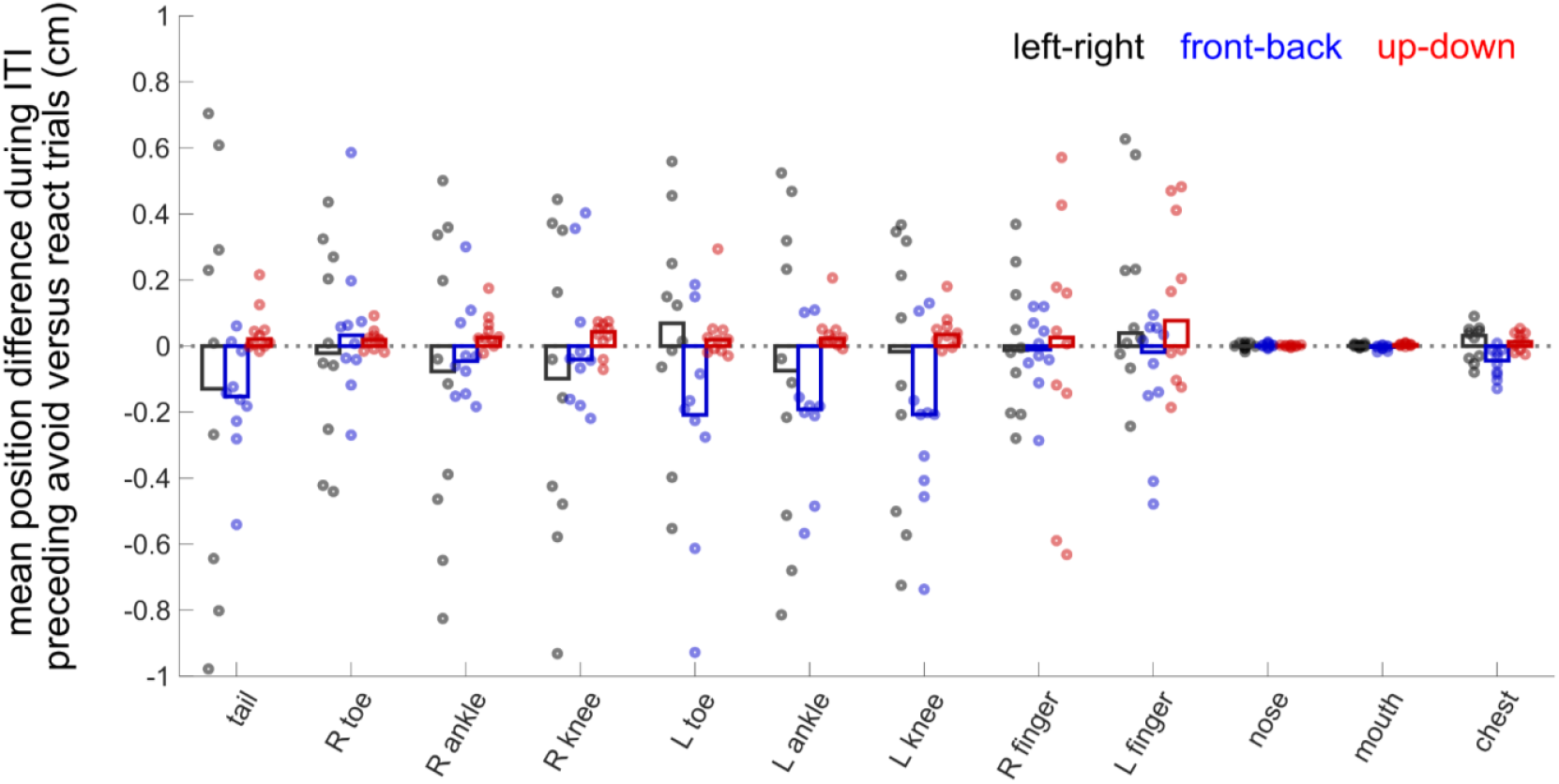
Idiosyncratic kinematic differences between avoid and react trials during the preceding intertrial interval (ITI). Mean differences (circles are individual mice, N=10; bars are mean across mice; colors represent the 3 Euclidean dimensions for each keypoint) in position between avoid and react trials, in the interval 1.5 sec before the cue, for all keypoints; the largest variation between mice is in the left-right direction (black), followed by front-back (blue), and up-down (red).

**Supplementary Figure 4:**
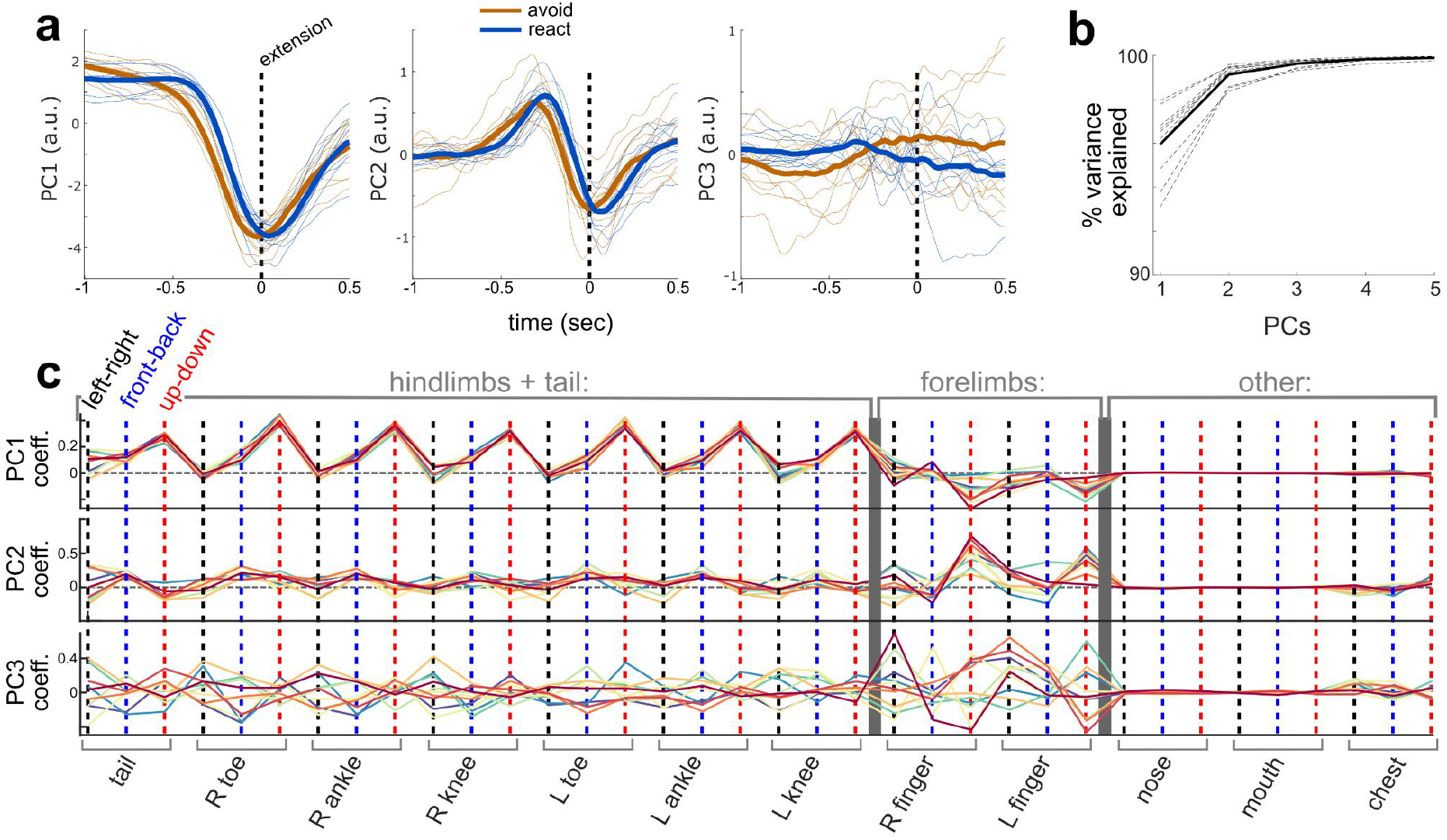
Kinematic state space from principal component analysis (PCA). **(a)** avoid (orange) and react (blue) trial mean projections onto the first 3 PCs of the kinematic state space derived from PCA using covariance of keypoints in the interval before extension threshold; thick lines are mean across mice, thin lines are individual mice avoid or react trial means projected onto PCs (N=10) **(b)** variance explained for the top kinematics PCs; dotted lines are individual mice, thick line is mean across mice (N=10) **(c)** PC coefficients for the first 3 PCs in kinematic state space, showing that the vast majority of movement variance (PC1) comes from up-down coordinates of hindlimb keypoints, consistently across all mice (different colored lines).

**Supplementary Figure 5:**
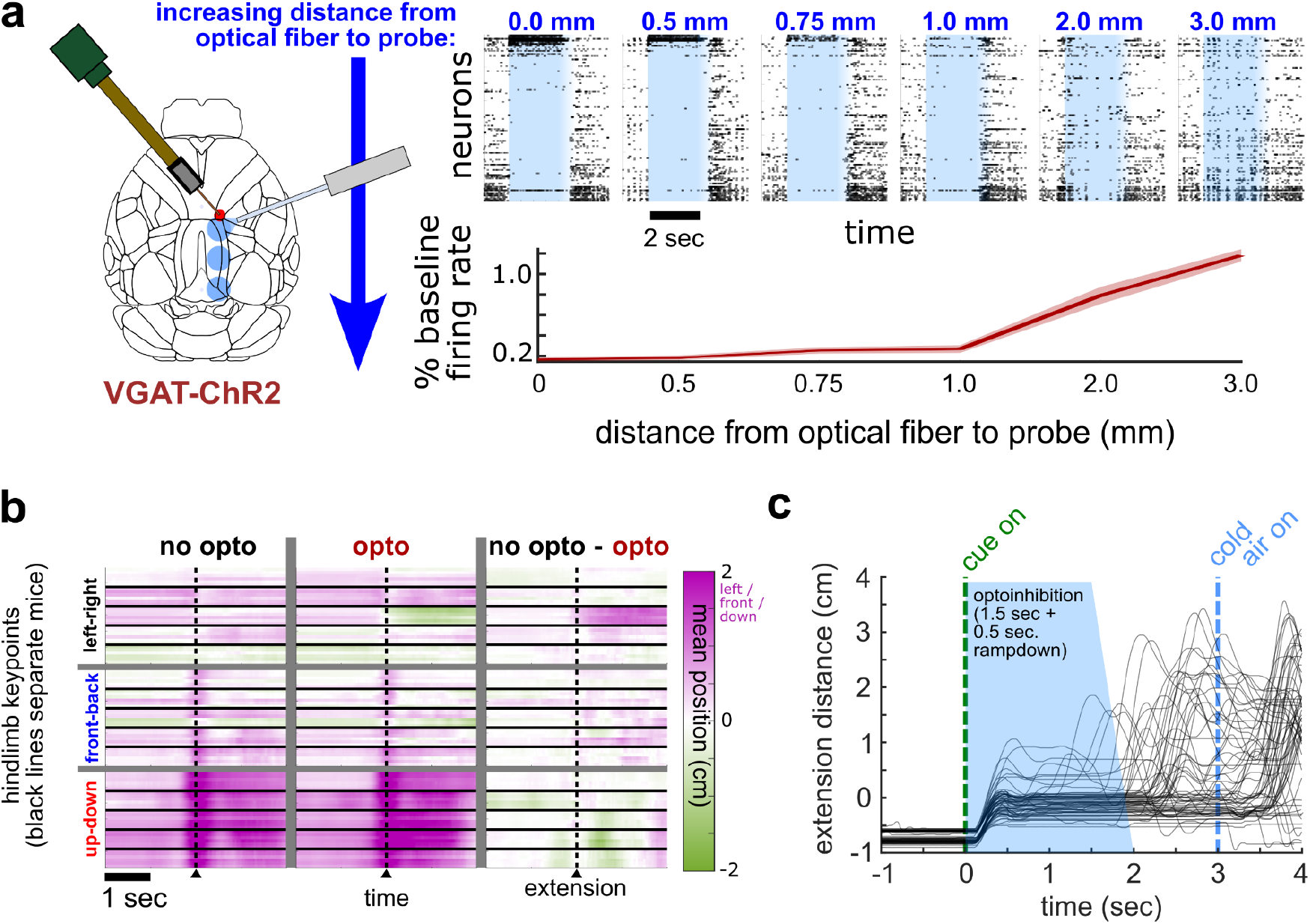
Local cortical optoinhibition details and controls. **(a)** local inhibition effect vs. distance from fiber (top, example mouse recording, bottom, mean ± s.e.m. N=3 mice) **(b)** heatmaps of mean position of all 6 hindlimb keypoints in interval around extension, with respect to the median resting ITI position, during a single session. Thick black lines separate different mice (N=10 mice), and vertical grey lines separate columns that show means on non-inhibited (react) trials, optoinhibited trials, and the difference between them **(c)** early release from optoinhibition reveals no sustained paralyzation due to nonspecific effect (N=4 mice), as mice can extend 100s of milliseconds after optoinhibition rampdown.

**Supplementary Figure 6:**
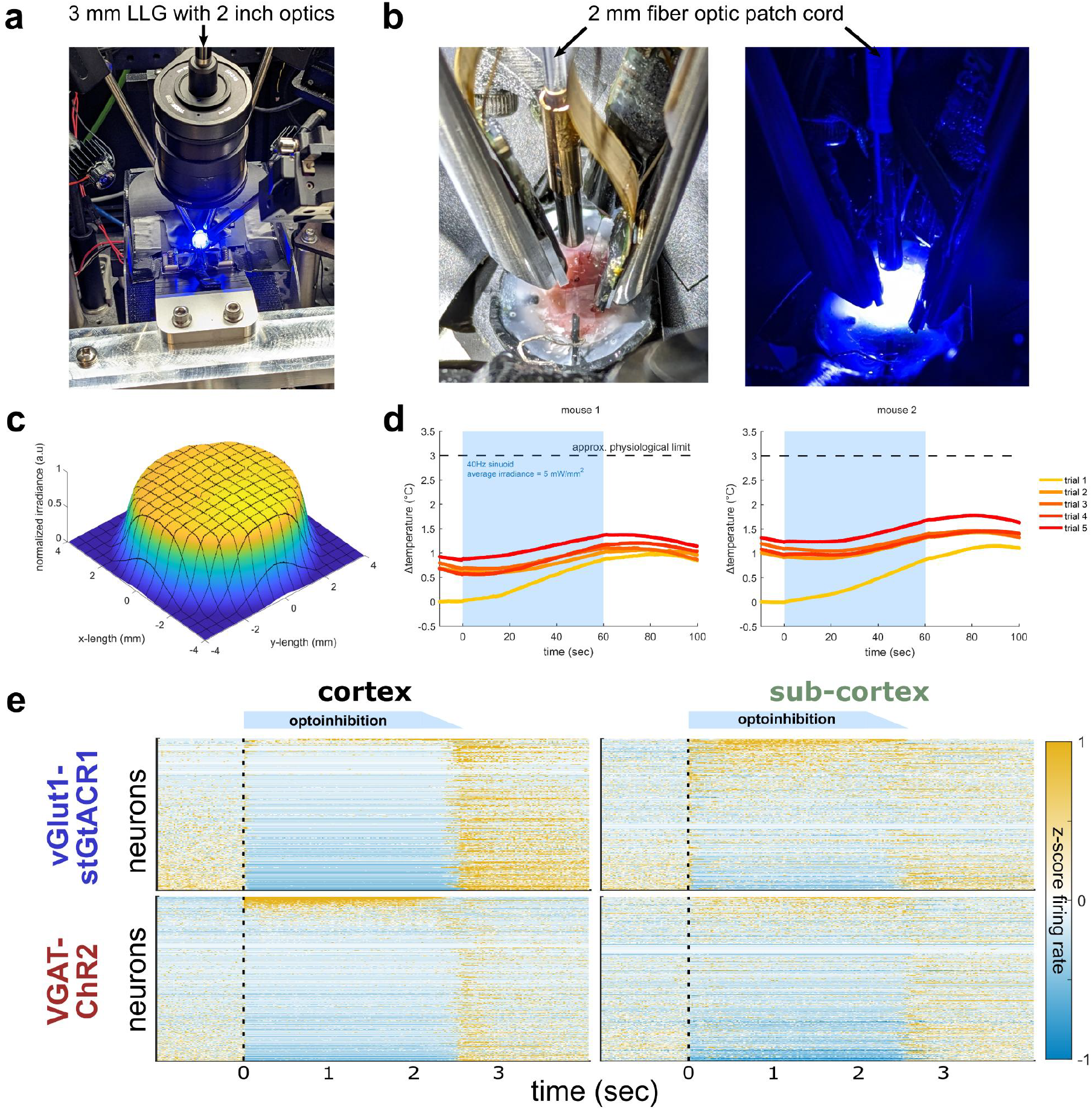
Acute decortication details. **(a)** photo of rig setup for liquid light guide (LLG) plus optics option for large scale inhibition (i.e. acute decortication) through the skull **(b)** photos of rig setup for 2mm fiber option for large scale inhibition through the skull, with 3 Neuropixels probes; right, before, and left, after LED light source turned on **(c)** measurement of light power spatial profile at the level of the skull **(d)** measurements of worst-case temperature increase at the skull, in the PBS bath continuous with cerebrospinal fluid, during five successive 1-minute, maximum power optoinhibition trials **(e)** comparison of mean responses to optoinhibition across all 3 probes (separated into cortical, left, and subcortical, right, neurons), in VGAT-ChR2 versus vGlut1-stGTACR1 mice; neurons are sorted by the magnitude of firing rate change during optoinhibition.

**Supplementary Figure 7:**
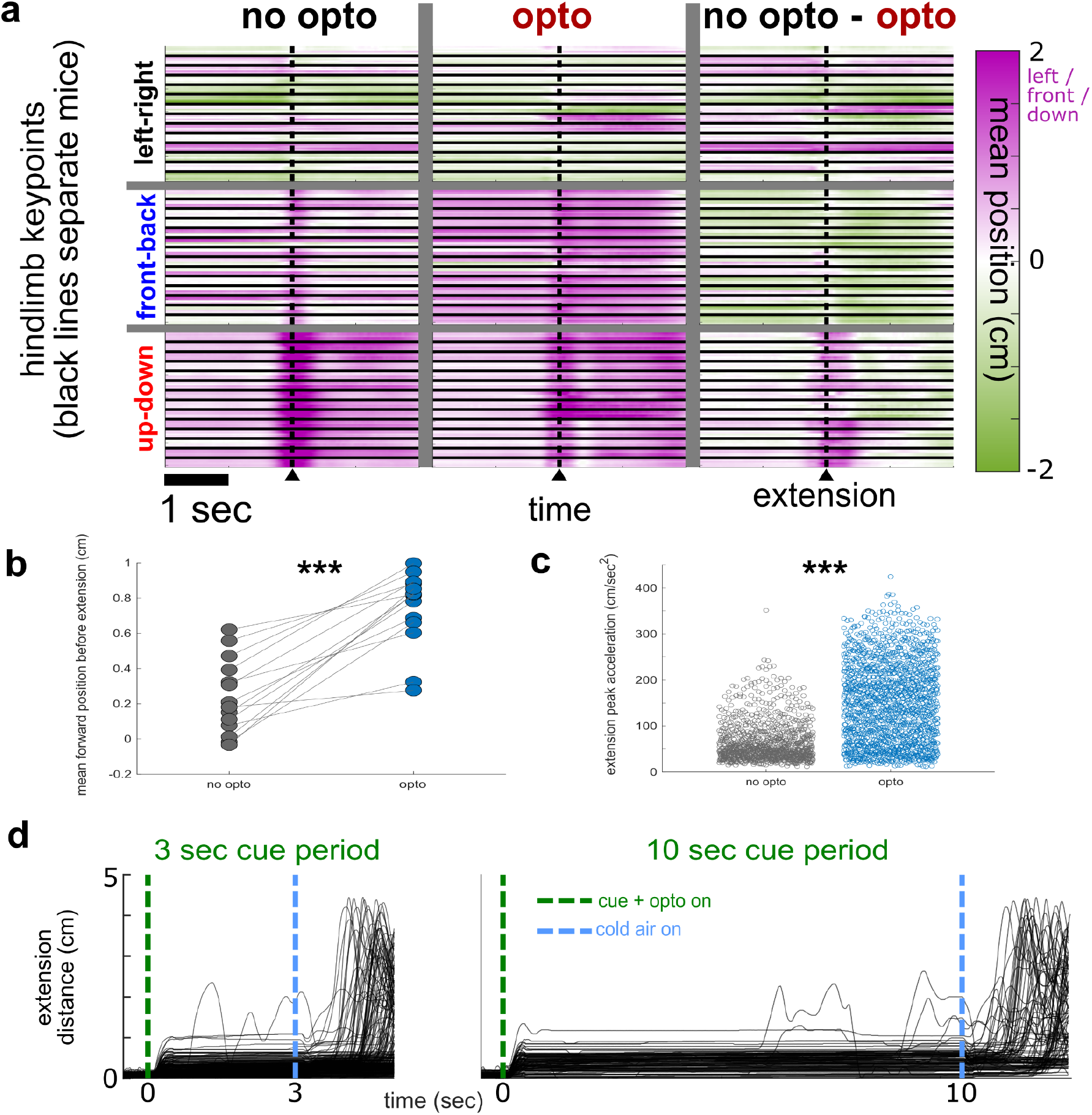
Acute decortication kinematics and controls. **(a)** heatmaps of mean position of all 6 hindlimb keypoints in interval around extension, with respect to the median resting ITI position, during a single session. Thick black lines separate different mice (N=14 mice), and vertical grey lines separate columns that show means on non-inhibited (react) trials, optoinhibited trials (having notable forward posture bias), and the difference between them **(b)** in mice with 3 days of acute decortication on 100% of trials, then release on day 4, non inhibited trials (including avoid trials) do not have the same forward posture bias, suggesting that detailed kinematic representations were not used for covert learning; p=1.22e^-12^ Wilcoxon signed rank test, N=14 mice **(c)** peak acceleration of platform extension for optoinhibited versus not trials, showing higher acceleration extensions during acute decortication; p=1.96e^-146^ Wilcoxon rank sum test, N=14 mice **(d)** additional experiments comparing extension distance (thin black lines are individual trials pooled across 6 mice) with 3 sec. cue period for acute decortication to a longer 10 sec. cue period, showing that mice generally do not extend even during a prolonged cue period (so behavioral effect of inhibition is not likely due to lack of preparation time).

**Supplementary Figure 8:**
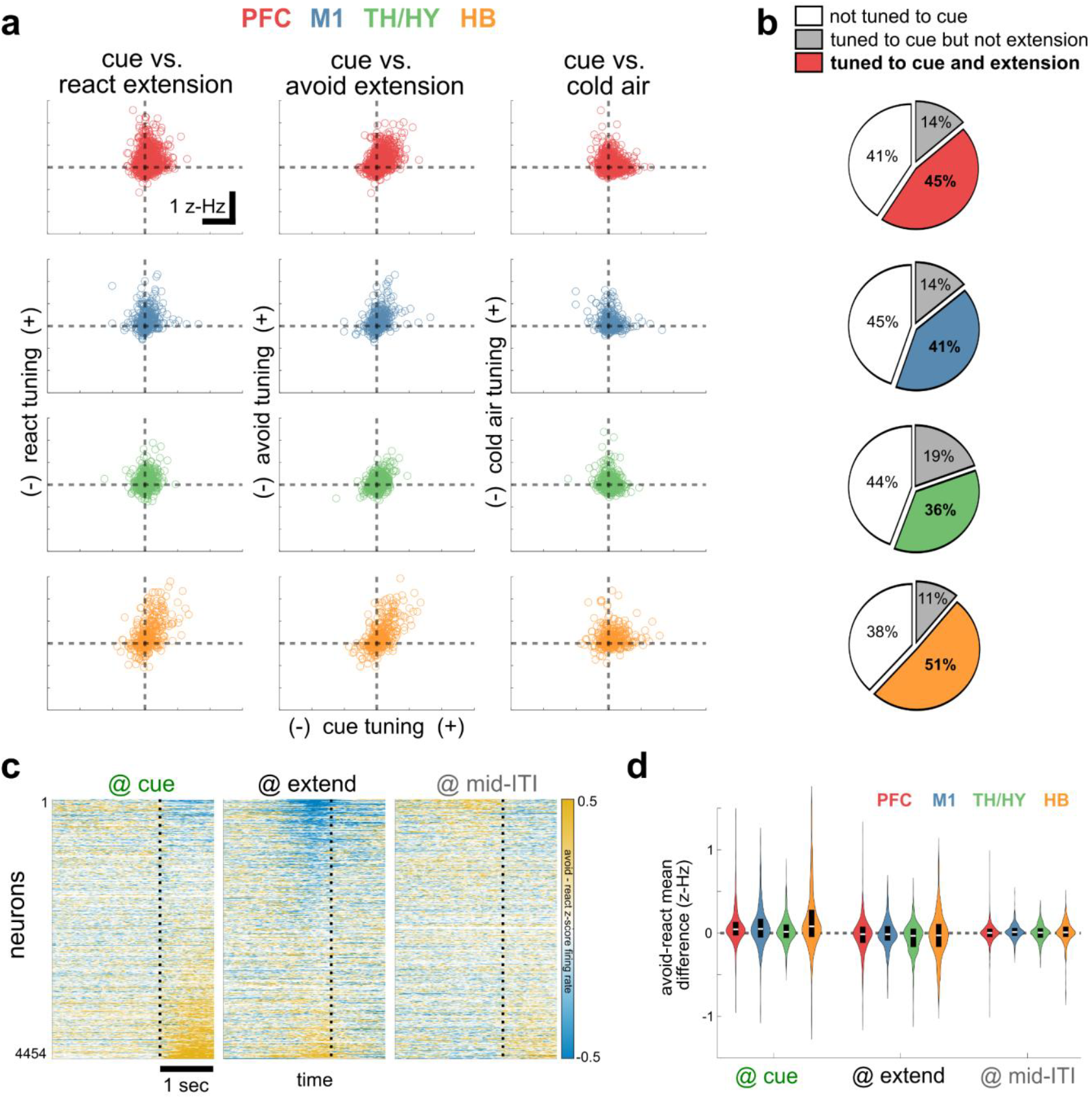
Widespread mixed-selective tuning of cue and movement and distributed modulations on avoid versus react trials. **(a)** comparison across broad brain areas of differential avoid-react tuning to cue (x-axis, mean z-score firing rate in 500 msec window after cue onset in avoid trials minus that in react trials), versus y-axis tuning to avoid or react extensions (mean z-score firing rate in 500 msec window before extension, relative to mid-ITI) or cold air (mean z-score firing rate in 100 msec window after cold air onset, relative to mid-ITI); open circles are single neurons, combined across 10 mice **(b)** pie charts for different brain areas showing percentage of neurons positively tuned to cue on avoid trials (non-white slices), and the percentage of those neurons also tuned to avoid extension (colored slices), which are the majority of non-white slices in all areas **(c)** mean z-score firing rate differences during avoid and react trials, combined across mice and brain areas and aligned at cue, extension, and mid-ITI (control); each heatmap is sorted separately by magnitude of modulation at aligned event **(d)** violin plots showing distributions of neurons in (c) separated into broad brain areas; colored violin area is proportional to full distribution, black boxes show middle half and white median.

**Supplementary Figure 9:**
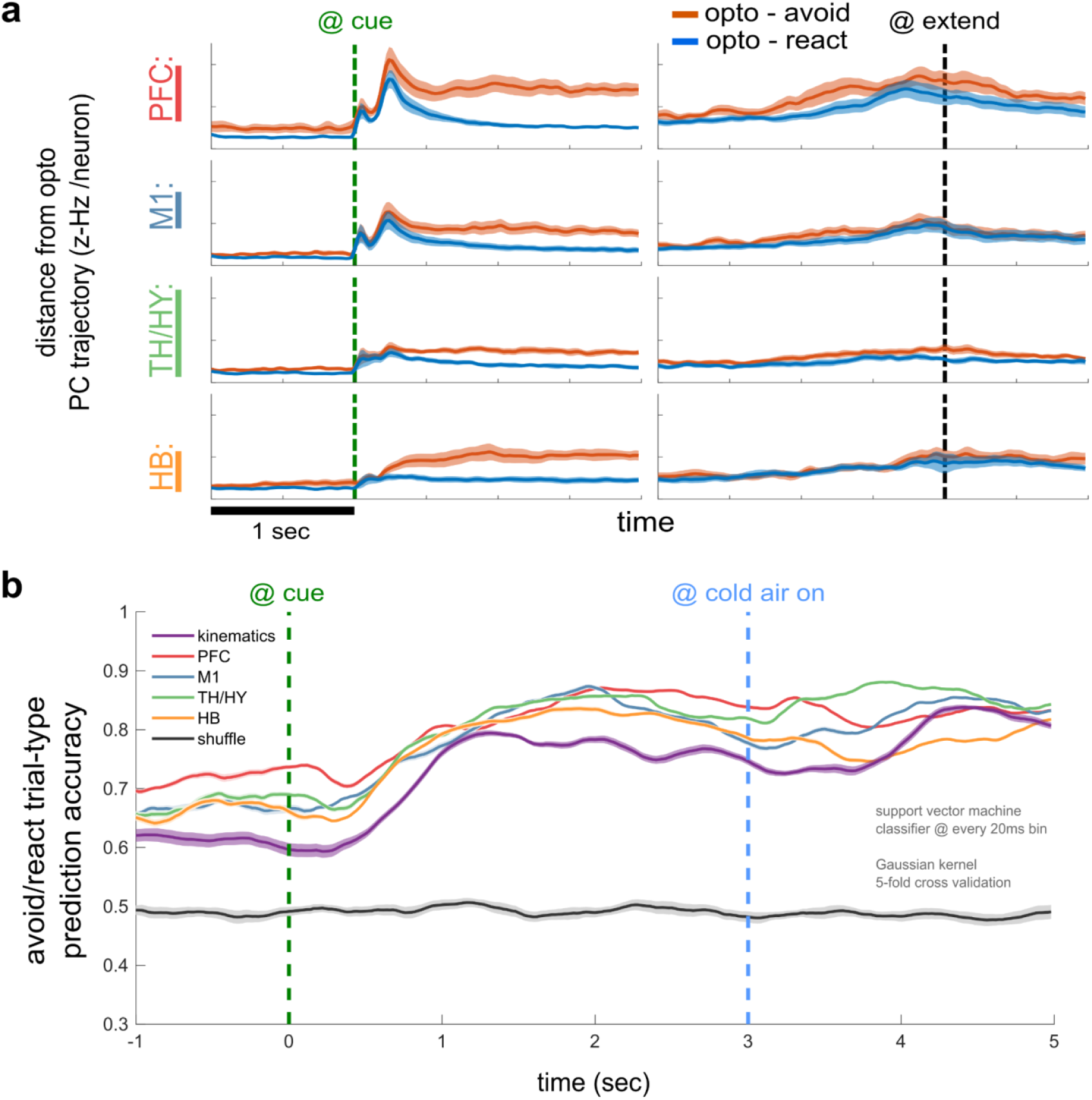
Cortical inhibition pushes neural activity towards react trajectories, and trial type can be decoded across the brain. **(a)** for different brain areas, mean ± s.e.m. PC distances between trajectories on optoinhibited trials versus either avoid (orange) or react (blue) trajectories, aligned to cue (left) and extension (right); activity on optoinhibited trials is closer to react trials in all cases **(b)** time-dependent trial type classifier showing mean ± s.e.m. prediction accuracy (N=10 trial sampling repeats) for different brain areas plus kinematics and shuffle controls; classification is done at each 20 msec time bin (aligned to cue) from trial data projected onto the top ten PCs for each area (see Methods).

## Supplementary Videos

**Supplementary Video 1: Interleaved avoid and react hindlimb extensions**. Ten example trial segments from a single mouse and session, showing mid-session avoid and react trials interleaved (orange ‘avoid’ or blue ‘react’ text denotes trial type). As in Figure 1, colored dots are automatically tracked keypoints, with teal lines added between them for clarity. Each trial segment is aligned to extension threshold, showing 500 msec before to 100 msec after. Playback speed is 1/4x real-time.

**Supplementary Video 2: Local frontal cortical optoinhibition disrupts avoid but not react hindlimb extensions**. Four consecutive trials from a mouse showing local optoinhibition effect on avoid extensions. Annotations in bottom left corner (‘beep’, ‘opto’, and ‘cold air’) denote when the auditory piezo cue, optoinhibition, and cold air are active. The trial sequence is (1) avoid, (2) optoinhibition / react, (3) avoid, (4) optoinhibition / react, without any ITI frames trimmed in between. Note that the mouse still extends almost immediately to the cold air during optoinhibition, but not to the cue / beep. Playback speed is real-time.

**Supplementary Video 3: Acute decortication can lead to forward posture during head-fixed hindlimb extension**. Single trial example of acute decortication in a VGAT-ChR2 mouse. Trial extension latency is 12.5 sec., and several attempts at extension are made in a forward posture before finally directing the extension downward. Playback speed is real-time.

